# Unlocking Bacterial Cellulose Functionalisation: Comparative Genetic and Co-Culture Strategies in *Komagataeibacter*

**DOI:** 10.64898/2026.01.12.699149

**Authors:** Anastasiya Kishkevich, Katie Gilmour, Maria Tsampika Manoli, M. Auxiliadora Prieto, Martyn Dade-Robertson, Meng Zhang, Tom Ellis

## Abstract

Bacterial cellulose (BC) is produced by diverse bacterial species, including members of the genus *Komagataeibacter*, and has emerged as a potential sustainable alternative to conventional materials such as plastics and leather. Its production by bacteria is rapid and easily scalable, and via genetic modifications or culturing the bacteria with other microbes, it is possible to rationally enhance the BC that is produced, for example by adding functional components. Here, we systematically evaluated BC functionalization strategies, including genetic engineering and co-culturing with other engineered microbes across the most widely used *Komagataeibacter* species: *K. rhaeticus*, *K. xylinus*, *K. medellinensis* and *K. sucrofermentans*. We established that all tested strains are amenable to DNA transformation and capable of expressing heterologous genes from identical genetic constructs. However, we identified species-specific differences in heterologous gene expression, cellulose production in varying environmental conditions and in the abilities of the strains to co-culture with *Escherichia coli*. Additionally, we demonstrated that all these cellulose-producing bacteria can establish functional symbiotic co-cultures with yeast. While inter-species variations in heterologous gene expression and co-culture dynamics are evident, BC modification through genetic engineering and co-culturing strategies remains achievable across all tested *Komagataeibacter* species. Our work provides a comparative framework to guide researchers in selecting optimal species based on their specific application requirements.

## Introduction

Bacterial cellulose (BC) has emerged as a versatile biomaterial, offering significant advantages over plant-derived cellulose. Biocompatibility and biodegradability combined with the three-dimensional network structure of BC makes it a promising candidate for applications ranging from biomedical devices and therapeutics to sustainable packaging materials and textiles. Numerous reviews have explored current and potential applications of bacterial cellulose ^1–14^.

A significant potential of BC lies in its programmable properties, made possible by the bacteria being open to genetic engineering and culturing with other microbes. Programmability allows the encoding of unique material properties at the stage of biosynthesis of material, in contrast to conventional post-production modification approaches. While BC is synthetised by many different bacteria species ^15,16^, members of *Komagataeibacter* genus are among the species with the highest cellulose yields and gaining increasing popularity in biotechnology, especially in the topic of Engineered Living Materials (ELMs) ^3,15,17–19^.

Genetic engineering of cellulose producing bacteria offers one approach to customise the properties and function of BC materials ^20–24^. To enable this engineering, sets of modular DNA parts, encoding constitutive and inducible promoters and Ribosome Binding Sites (RBS) are available for programming the expression of heterologous proteins in *Komagataeibacter rhaeticus, K. xylinus* and *Novacetimonas hansenii* ^20,22,23,25^. The *Komagataeibacter* Toolkit (KTK) system developed by Goosens *et al*. offers a modular Golden Gate based approach to design and construct multigene expression cassettes for *K. rhaeticus*, as well as genome integration cassettes too ^23^. In past work, KTK has been used to produce colourful BC when cellulose-producing bacteria are directed to express red fluorescence proteins and chromoproteins ^20,23^. It has also been used to engineer BC-producing bacteria to express the tyrosinase enzyme to convert tyrosine into melanin, leading to the growth of colourfast black BC materials as a leather alternative ^24^. Genome engineering of *Komagataeibacter* is also possible. CRISPR-Cas9-mediated gene deletions, directed evolution and targeted gene disruption have all been successfully employed in studies for altering BC properties and yield _22,23,26–31._

While there is increasing progress in engineering cellulose-producing bacteria, organisms such as *Escherichia coli* or baker’s yeast *Saccharomyces cerevisiae* are much more established as engineerable hosts for producing functional molecules and proteins. Co-culturing BC-producing bacteria with these model microbes offers a promising trajectory for functionalisation of BC. Engineered *E. coli* can be added to already-formed BC pellicles and will be entrapped forming programmable living material ^32–35^. A similar approach has been demonstrated with *S. cerevisiae,* immobilising living yeast within sterile pellicles ^36^. In more advanced work, a responsive living material was developed from Synthetic Symbiotic Co-cultures Of Bacteria and Yeast (SynSCOBY) where *K. rhaeticus* was co-cultured with engineered yeast from the very start of pellicle formation^37^. Yeast were engineered to either secrete enzymes that incorporate into growing material or to express reporter proteins in response to a chemical inducer signal. Indeed*, de novo* co-culturing has also been successful for *Komagataeibacter* partnered with many other microbes, including *Lactobacillus, Leuconostoc, Xanthomonas, Pseudomonas, Ralstonia* and *Bacillus* ^38–47^.

Here, we evaluated four widely-studied *Komagataeibacter* species (*K. rhaeticus*, *K. xylinus*, *K. medellinensis* and *K. sucrofermentans* ^20,27,48,49^) for their potential in engineered living material research, by comparing their cellulose production, genetic tractability and suitability for co-culturing. Working with five strains representative of commonly used or industrially optimised versions of the four species, we demonstrated that the KTK system and plasmids originally developed for *K. rhaeticus* iGEM are broadly applicable for genetic engineering across the *Komagataeibacter* species and can be used to heterologously express proteins that visibly change the material. We further established that the *Komagataeibacter* strains can form productive co-cultures with engineered *E. coli* and *S. cerevisiae* yeast strains to produce programmable living materials, with both microbes successfully integrating into pellicles produced by nearly all the tested *Komagataeibacter* species. Together, this work provides a comparative framework to guide researchers in selecting the optimal cellulose-producing bacterial strains for specific ELM applications.

## Results

### The Komagataeibacter Toolkit is functional across Komagataeibacter

The KTK offers a modular molecular cloning platform for genetic engineering and heterologous protein expression in *Komagataeibacter* species ^23^. Originally developed for the *K. rhaeticus* iGEM strain, we sought to determine whether KTK expression cassettes could function across a broader spectrum of *Komagataeibacter* species commonly employed in research and biotechnology. To assess this we focused on five strains, four that are representative of the most commonly-studied species of *Komagataeibacter* (*K. rhaeticus* iGEM, *K. xylinus* DSM-2325, *K. sucrofermentans* DSM-15973, *K. medellinensis* CECT 8140 ID13488) ^50^ and a strain evolved for industrial applications (*K. medellinensis* CIB-4 from the Prieto group) ^49^.

To test cross-species compatibility of KTK, we first assembled a plasmid expressing the violet-blue chromoprotein AmilCP (KTK_448) and took a previously-made plasmid carrying the purple chromoprotein gfasPurple (PurpleCP, KTK_349) ^23^ and electroporated both into competent *Komagataeibacter* cells, selecting for growth on antibiotic-containing plates. Transformed colonies displayed striking visual differences, expressing various shades of purple (PurpleCP) and violet-blue (AmilCP) depending on the strain used (**Fig. 1A**). The industrial strain *K. medellinensis* CIB-4 exhibited the most vibrant coloration for both chromoproteins, while *K. rhaeticus* and *K. sucrofermentans* colonies produced nearly indistinguishable hues. *K. medellinensis* CECT 8140 showed intermediate intensity, brighter than *K. rhaeticus* but less vivid than the CIB-4 strain. Notably, *K. xylinus* displayed an intriguing pattern: PurpleCP expression appeared less intense than in *K. rhaeticus* and *K. sucrofermentans*, whereas AmilCP expression was markedly stronger. Strain-specific variation in chromoprotein expression extended to the cellulose pellicles produced by these bacteria (**Fig. 1B**). *K. medellinensis* CIB-4 again generated the most intense colour, while the expression patterns in *K. xylinus*, weaker PurpleCP but stronger AmilCP, mirrored the colony phenotypes. Pellicles from *K. medellinensis* CECT 8140 exhibited coloration comparable to *K. rhaeticus*.

**Figure 1.**
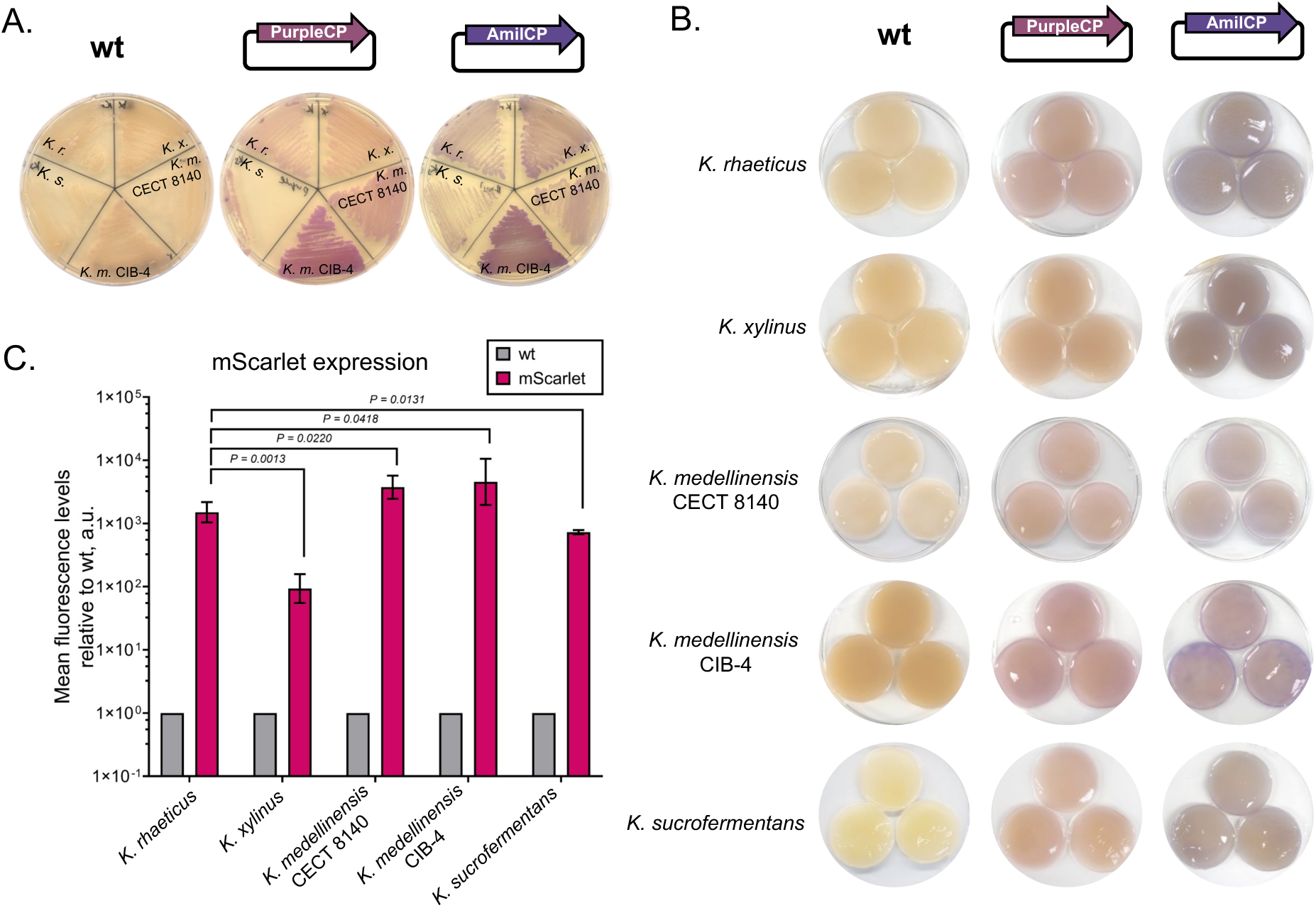
Constitutive expression of chromoproteins and red fluorescent protein in different *Komagataeibacter spp*. **A.** Bacteria cells expressing chromoproteins from KTK_349 (PurpleCP) and KTK_448 (AmilCP) grown on YPD-Chloramphenicol plates, wt strains grown on YPD. *K.r. = K. rhaeticus, K.x. = K. xylinus, K.m. CECT 8140 = K. medellinensis CECT 8140, K.m.CIB-4 = K. medellinensis CIB-4, K.s. = K. sucrofermentas.* **B.** Bacterial cellulose formed at 30 °C from cells expressing chromoproteins. **C.** Expression of RFP from KTK_292 plasmid (mScarlet) in all strains analysed by flow cytometry, mean fluorescence values are normalised to the wildtype (wt) values, n=5 for mScarlet strains, n=3 for wt strains, error bars denote standard deviation; data was analysed with GraphPad using unpaired t-test with Welch’s correlation.

To quantitatively assess heterologous protein expression across strains, we transformed all five strains with plasmid KTK_292 expressing the red fluorescent protein mScarlet ^23^ and analysed fluorescence intensity in living cells via flow cytometry (**Fig. 1C**). All tested strains demonstrated elevated red fluorescence relative to untransformed controls, confirming successful expression. Both *K. medellinensis* strains, achieved significantly higher fluorescence levels than *K. rhaeticus*. Conversely, *K. xylinus* and *K. sucrofermentans* exhibited lower expression, with *K. xylinus* fluorescence approximately 15-fold less compared to that of *K. rhaeticus*.

Collectively, these results establish that KTK-based plasmids can be successfully transformed into diverse *Komagataeibacter* species and drive heterologous protein expression across phylogenetically distinct strains, albeit with substantial strain-dependent variation in expression levels.

Building on our chromoprotein screening results, which revealed AmilCP’s superior visibility in pellicles, we selected this violet-blue reporter for subsequent experiments. To expand beyond constitutive expression, we leveraged the inducible gene expression system in KTK that is based on acyl-homoserine lactone (AHL) signalling. We constructed dual-cassette plasmids containing two transcriptional units: LuxR expressed from the constitutive J23104 promoter and AmilCP controlled by the AHL-responsive pLux promoter. In this quorum-sensing circuit, AHL binding to LuxR inside cells triggers conformational changes that activate transcription from the pLux promoter (Fig. 2A).

**Figure 2.**
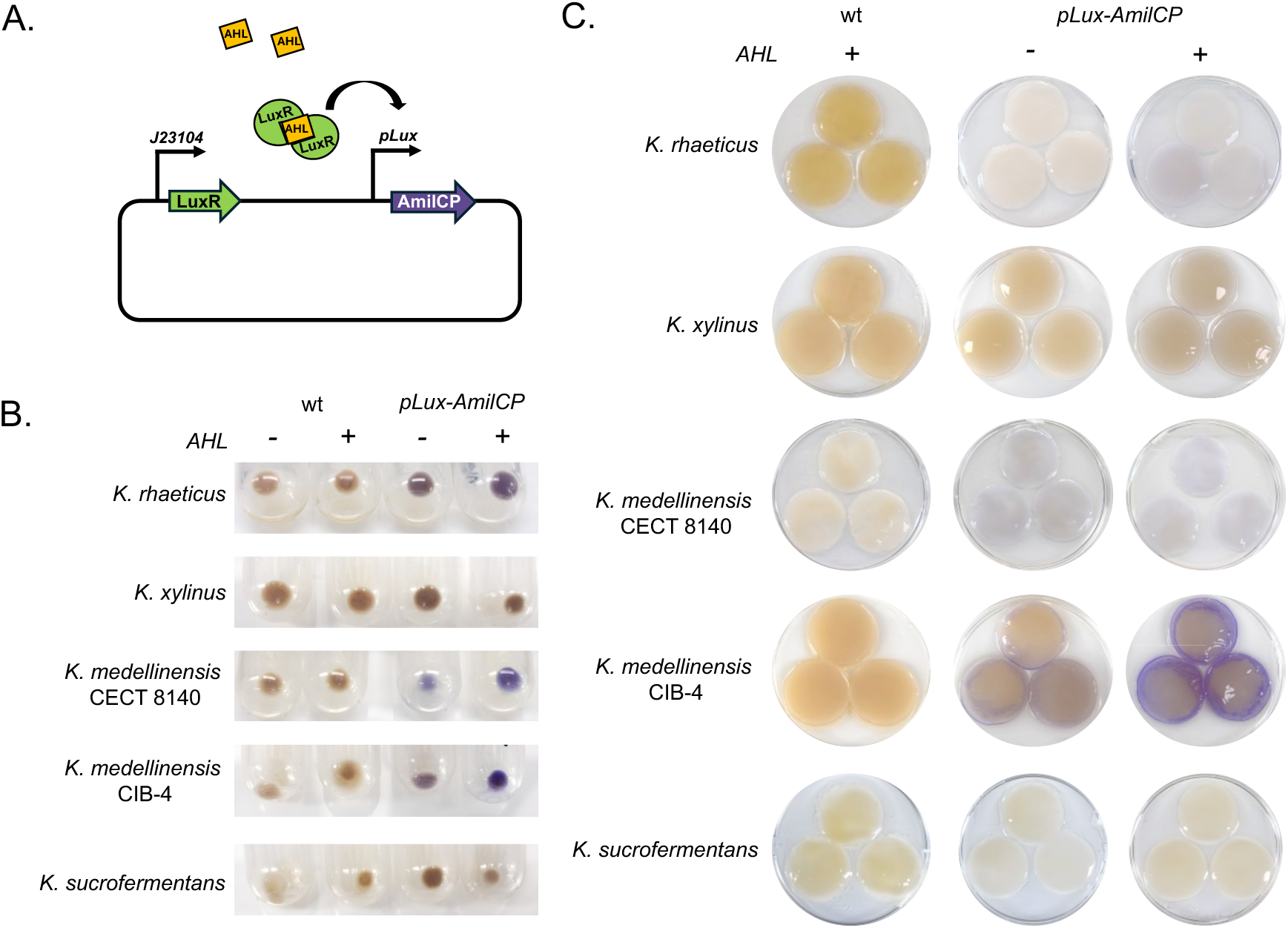
Induction of AmilCP expression in different *Komagataeibacter spp*. **A.** Diagram of AHL inducible genetic circuit. AHL is externally added and diffuses into the cell. LuxR binds AHL and forms a complex that activates transcription from pLux promoter. **B**. Cell pellets from cultures (wt and cells with the plasmid construct) grown with and without 1 µM AHL. Cultures were grown to high density, divided into 2 tubes and grown with and without AHL. Basal AmilCP expression is observed in non-induced conditions. **C.** BC pellicles produced by wt and plasmid-containing bacteria. Pellicles were grown at 30°C for 3-4 days prior addition to AHL and left for 3 more days to express AmilCP. wt pellicles with AHL were used as control to show that 1 µM AHL (in DMSO) does not disrupt pellicle integrity.

Cells were grown for 72 hours in liquid culture before 72-hour induction with AHL. After induction we observed markedly different responses across *Komagataeibacter* strains (Fig. 2B). While uninduced cultures exhibited basal AmilCP expression, *K. rhaeticus* and both *K. medellinensis* strains (CECT 8140 and CIB-4) displayed pronounced colour intensification upon AHL addition. But strikingly, *K. xylinus* and *K. sucrofermentans* showed no detectable AmilCP expression. In this AHL-inducible genetic construct we used identical promoter-RBS combinations to drive LuxR expression as were used for constitutive RFP expression in Fig. 1C. Notably in that data, *K. xylinus* and *K. sucrofermentans* showed lower expression that the other strains. Therefore, we attribute the lack of AmilCP induction in these strains as a result of insufficient LuxR expression. Alternatively, is also possible that transcriptional machinery in *K. xylinus* and *K. sucrofermentans* does not drive transcription from the pLux promoter. The inducible expression patterns manifested visibly in bacterial cellulose (BC) pellicles (Fig. 2C).

*K. medellinensis* CIB-4 generated the most vividly coloured pellicles, featuring an exceptionally bright violet-blue rim. This spatial pattern likely reflects oxygen gradients or metabolic zonation during pellicle formation. *K. rhaeticus* and *K. medellinensis* CECT 8140 produced BC with subtle violet-blue tinting throughout. Intriguingly, while *K. xylinus* cells in liquid culture showed no AmilCP expression, their BC pellicles developed a faint blue rim. *K. sucrofermentans* pellicles remained colourless, consistent with the absence of expression seen in cell pellets. Overall, pellicle coloration intensity correlated closely with cellular AmilCP levels.

As before, this work demonstrated that the same KTK plasmid constructs function across diverse *Komagataeibacter species*, but genetic and physiological differences among strains cause variation in heterologous gene expression.

### Co-culturing *Komagataeibacter* with other bacteria

Co-culturing *Komagataeibacter* with other microbes offers an alternative strategy for functionalising BC and modifying its material properties. Previous studies demonstrated that co-cultures of cellulose-producing bacteria with *E. coli* can generate functionalised pellicles ^39,40^. *E. coli* is a convenient chassis as it has minimal culture requirements and genetic engineering is well-studied. We designed LacI-inducible T7 promoter expression cassettes for *E. coli* BL21 to produce the chromoproteins AmilCP, RedCP and the fluorescent protein, GFP. In the case of the GFP and RedCP constructs, the genes were both designed to encode fusion proteins so that the proteins were fused to synthetic cellulose-binding motifs (‘double-CBM’ and CBMcex) ^51,52^, enabling the binding of the fusion protein to cellulose fibres if it can be released from the *E. coli* cell (e.g. during natural cell lysis).

The three engineered *E. coli* and *E. coli* carrying an empty plasmid were co-cultured in a pairwise fashion with the five *Komagataeibacter* strains in rich medium at room temperature (22 °C) for 14 days. This temperature was chosen as it is suitable for stable T7 promoter-based expression in *E. coli* and is also a permissible temperature for BC growth. IPTG induction was delayed 24 hours post-inoculation to allow stable *E. coli* population establishment before imposing the burden of heterologous protein expression. Following pellicle formation, samples were stored at 4 °C to aid complete chromoprotein maturation.

The 20 co-cultures yielded a heterogeneous spectrum of functionalised pellicles (Fig. 3). Notably, all *K. medellinensis* CECT 8140 co-cultures demonstrated no detectable *E. coli* incorporation. However, *E. coli* incorporation was seen in the other four strains. Presence of AmilCP was barely detectable in all cases, with only *K. xylinus* pellicles displaying a visibly faint blue tint. However, GFP-CBM and RedCP-CBM were easily detectable in pellicles under UV illumination. The strength and uniformity of protein signal varied between strains, with *K. xylinus - E. coli* co-cultures, generally giving the best results. Incomplete colouration of the pellicles by the *E. coli* was observed when co-culturing with the *K. rhaeticus* and *K. sucrofermentans* strains.

**Figure 3.**
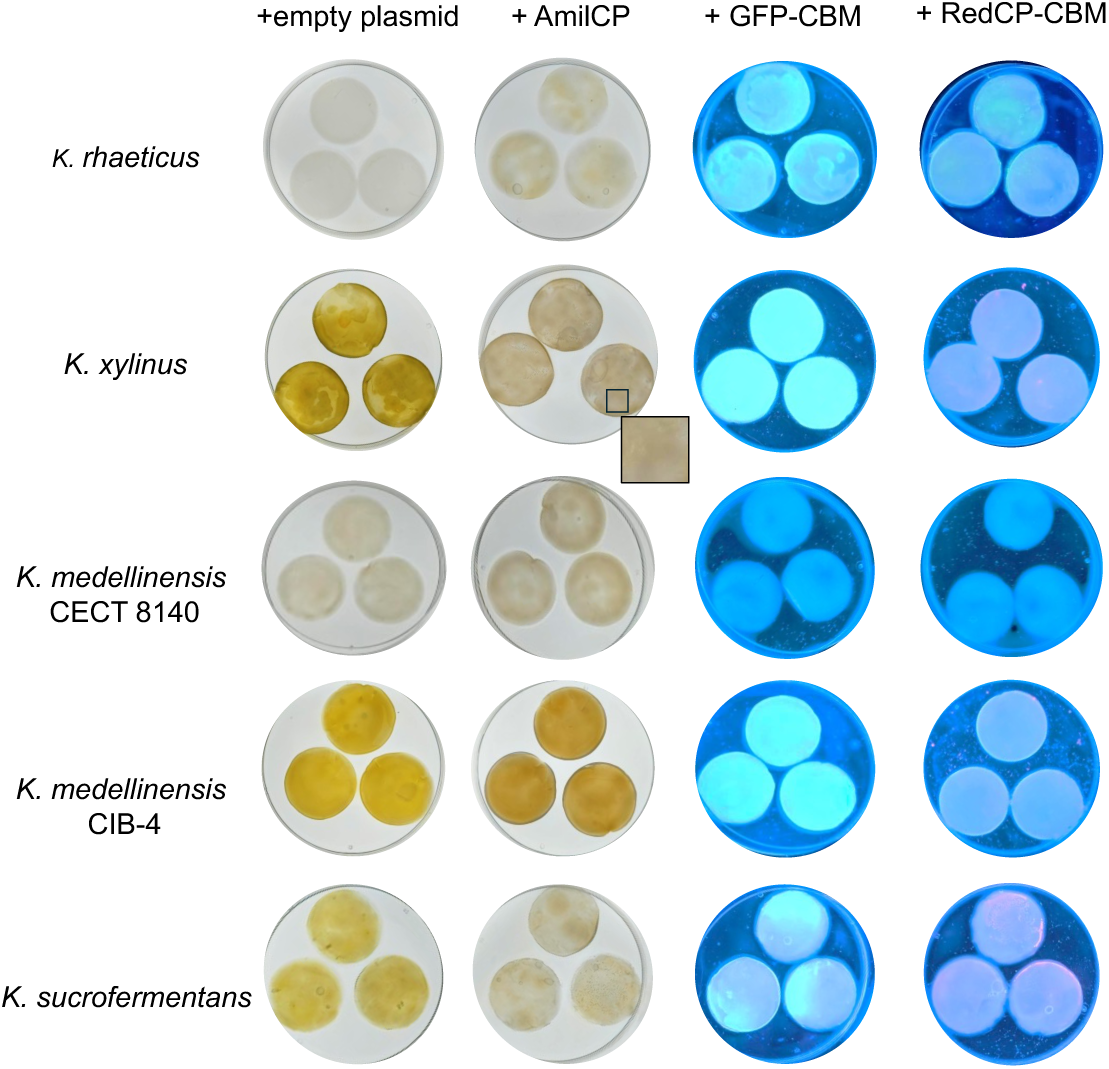
Functionalisation of bacterial cellulose from co-culturing with engineered *E. coli*. Cellulose-producing bacteria and *E. coli* co-cultures expressing visibly detectable proteins were grown for 14 days at room temperature. *E. coli* carried either empty plasmid, AmilCP, GFP-CBM or RedCP-CBM. Pellicles were kept at 4 °C prior imaging to allow chromoproteins to mature. Images were taken with room lights (empty plasmid and AmilCP) or UV lights at 254/312 nm (GFP-CBM and RedCP-CBM).

To enhance programmable material properties, we next established a quorum sensing-based intrakingdom communication system where *E. coli* functions as the signal sender to induce gene expression in *Komagataeibacter* receiver cells. For this we used previously developed AHL-mediated communication plasmids ^20,21^ with *E. coli* constitutively expressing the AHL synthase LuxI (sender), triggering AmilCP expression in BC-producing cells (receiver) (Fig. 4A). Co-cultures were established as before and maintained for 14 days at room temperature. Pellicles treated with commercial AHL served as positive controls for AmilCP induction.

**Figure 4.**
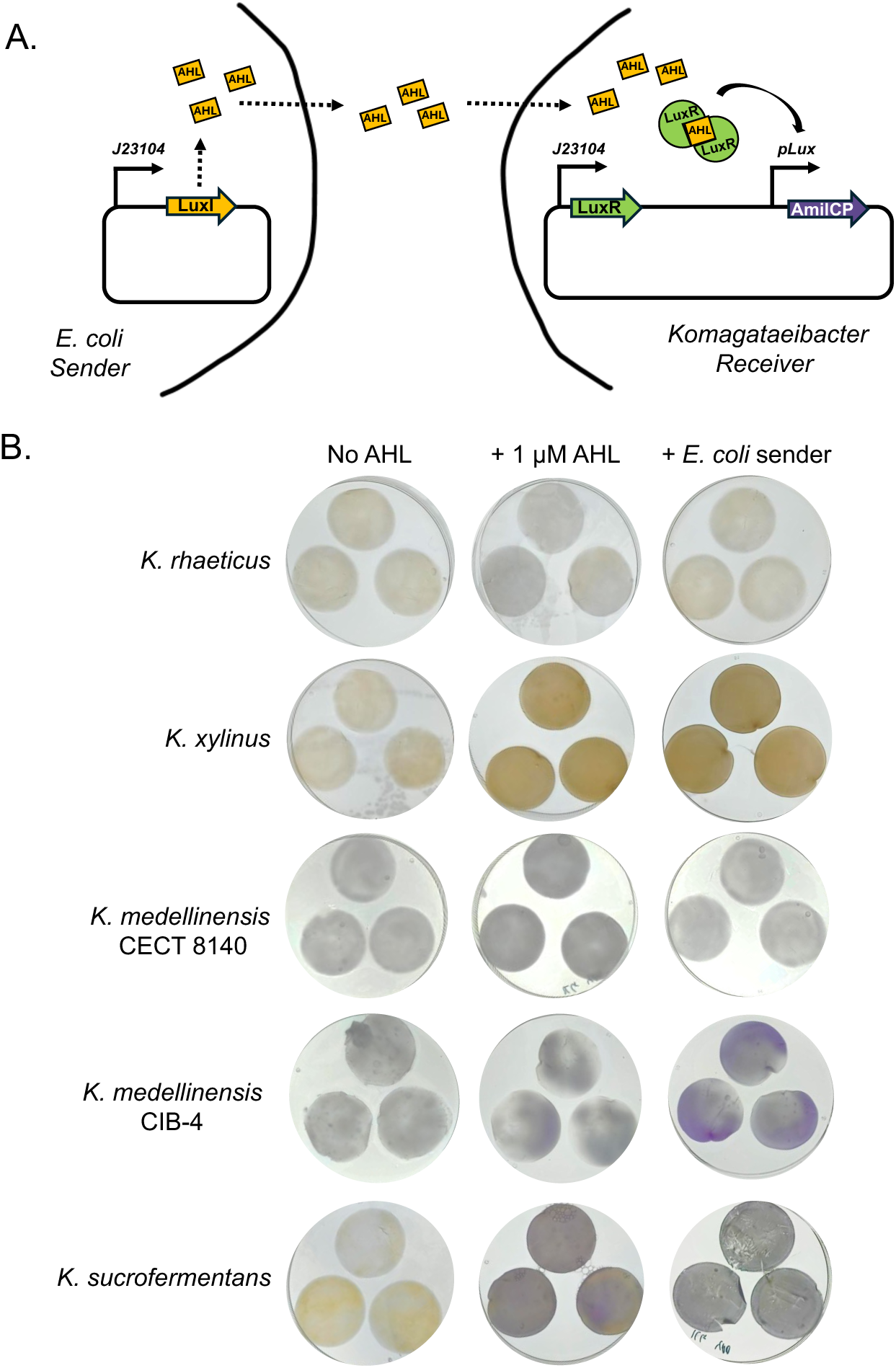
Functionalisation of bacterial cellulose via communication between engineered *E. coli* and cellulose producing bacteria. A. Diagram of cell-to-cell communication between bacteria in co-culture. *E. coli* expresses acyl-homoserine lactone synthase LuxI. AHL produced by *E. coli* enters *Komagataeibacter* cells; LuxR binds AHL and activates AmilCP transcription from pLux promoter. **B**. BC pellicles produced by cellulose-producing bacteria expressing AmilCP under pLux promoter and *E. coli* constitutively expressing LuxR. Pellicles were grown at room temperature for 14 days and kept at 4°C prior imaging to allow chromoproteins to mature. Pellicles produced by monocultures and treated with 1 µM of AHL after 24 hours are used for comparison.

The results once again showed variation in functionalization between the BC-producing strains (Fig. 4B). This time *K. xylinus* pellicles showed no coloration in either co-cultures or AHL-treated controls, whereas *K. medellinensis* CECT 8140 showed coloration in all cases, corresponding to the prior results in Fig. 3. The strongest colouration of BC was produced when co-culturing *E. coli* with *K. medellinensis* CIB-4, which intriguingly outperformed the visible AmilCP output from the same BC-producing bacteria when simply given commercial AHL.

Out of the five BC-producing strains, only *K. sucrofermentans* exhibited the pattern of expression that would be expected where BC grown in the control conditions is absent of AmilCP (2 out of 3 samples), but the AmilCP is visible when AHL is applied or the *E. coli* sender cells are present in co-culture. The expected pattern was partially seen with *K. rhaeticus* pellicles, however no AmilCP expression was evident from the attempted co-culture with *E. coli*. We hypothesised that in this case *K. rhaeticus* had failed to establish a productive co-culture with *E. coli* at room temperature.

To check this hypothesis, we next quantified *E. coli* incorporation into pellicles using flow cytometry. *E. coli* constitutively expressing GFP were co-cultured with untransformed *Komagataeibacter* strains as described above. GFP fluorescence was used to enable discrimination between the two types of bacteria which have similar sizes. The percentage of GFP-positive cells within cellulase -digested pellicles was calculated as the *E. coli* incorporation efficiency **(Fig. S1**).

Flow cytometry revealed substantial batch-to-batch variability across independent experiments (Fig. 5A). Pellicles produced by *K. xylinus*, *K. medellinensis* CIB-4, and *K. sucrofermentans* contained median *E. coli* populations of 17.12%, 18.34%, and 19.34% of the total cell counts, respectively. In contrast, *K. medellinensis* CECT 8140 and *K. rhaeticus* showed markedly divergent incorporation patterns. *K. rhaeticus* pellicles displayed pronounced inconsistency in *E. coli* incorporation (Fig. 5B) with some batches containing no detectable *E. coli*, while others reached 25-30% incorporation. Pellicles with low *E. coli* content were notably thinner.

**Figure 5.**
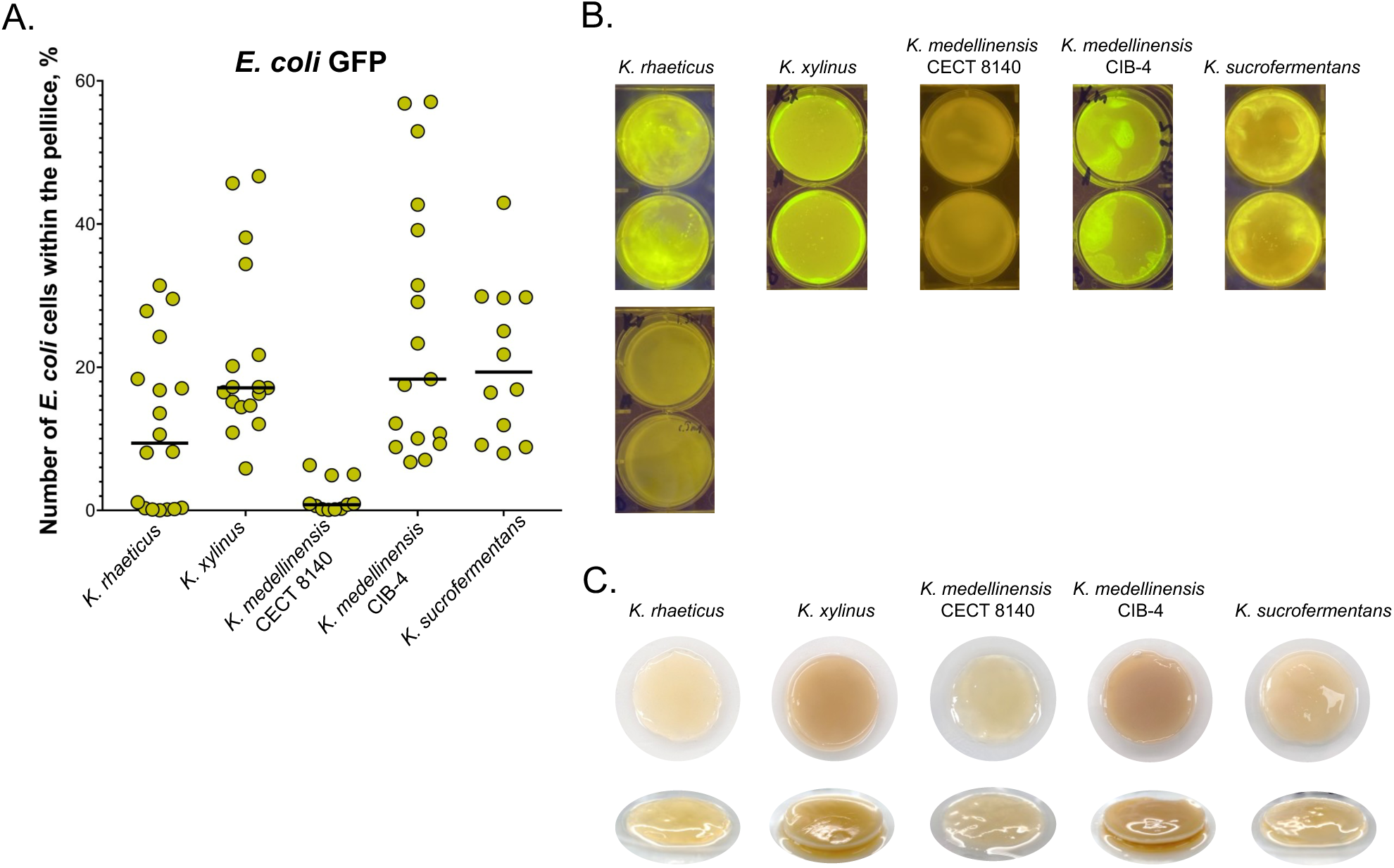
*Komagataeibacter* - *E. coli* co-cultures and pellicle production at room temperature. **A.** Analysis of *E. coli* incorporation into cellulose pellicles by flow cytometry. Pellicles produced by co-cultures of Komagataeibacter strains and *E. coli* constitutively expressing GFP were grown at room temperature for 14 days. Pellicles were digested in PBS+2% cellulase and 2 mL samples were analysed by flow cytometry. Data was processed with GraphPad (significant outliers removed) and % of GFP positive events within the analysed sample was plotted. **B.** Representative images of pellicles analysed by flow cytometry (under blue lights). Incorporation of *E. coli* is indicated by bright fluorescence. Two sets of pellicles for *K. rhaeticus* show variations between different experiments. **C.** Pellicles grown from monoculture of cellulose-producing bacteria at room temperature for 14 days.

We hypothesised that *Komagataeibacter* strains exhibit different growth performance at room temperature. Growth curve analysis at 22°C and 30°C revealed that *K. rhaeticus* and *K. medellinensis* CECT 8140 exhibited extended lag phases; 22-24 hours at 22°C compared to 10 hours at 30°C (**Fig. S2**). Nevertheless, all strains achieved comparable optical densities (OD_600nm_) after 100 hours of culture incubation. This growth behaviour correlated with monoculture pellicle formation at 22°C: strains with prolonged lag phases produced thinner, weaker pellicles (Fig. 5C), whereas the *K. xylinus* and *K. medellinensis* CIB-4 strains generated robust pellicles at room temperature comparable to those formed at 30°C. These findings indicate that room temperature represents suboptimal conditions for *K. rhaeticus* and *K. medellinensis* CECT 8140 pellicle formation and *E. coli* co-culture integration.

### Functionalisation of bacterial cellulose via SynSCOBY

Prior work developed the synthetic symbiotic co-culture of bacteria and yeast (SynSCOBY) to produce ELMs from engineered *S. cerevisiae* co-cultured with *K. rhaeticus* ^37^. We next sought to evaluate if this approach could work with the other *Komagataeibacter* strains. The original SynSCOBY formulation comprised 45% density gradient medium (Optiprep™) mixed with standard Yeast/Peptone/Sucrose medium, yielding a final 0.5X YPS concentration. To enhance nutrient availability, we increased YPS concentration to 1X in the final co-culture medium for this study.

SynSCOBYs were cultivated at room temperature (22 °C) and at 30 °C in both 0.5X and 1X YPS formulations. Yeast incorporation into pellicles was quantified by flow cytometry, with yeast cells distinguished from bacteria based on their distinct forward and side scatter profiles (**Fig. S3**). The percentage of yeast cells was calculated relative to total microbial cells (bacteria plus yeast) within digested BC pellicle samples. As observed in the above *E. coli* co-cultures, substantial batch-to-batch variability occurred across independent SynSCOBY experiments, presumably caused by heterogeneous nature of BC cultures (Fig. 6). 1X YPS medium at 30 °C yielded the highest yeast incorporation for all SynSCOBY combinations, except for *K. medellinensis* CIB-4, which showed comparable values at 22 °C and 30 °C in 1X YPS.

**Figure 6.**
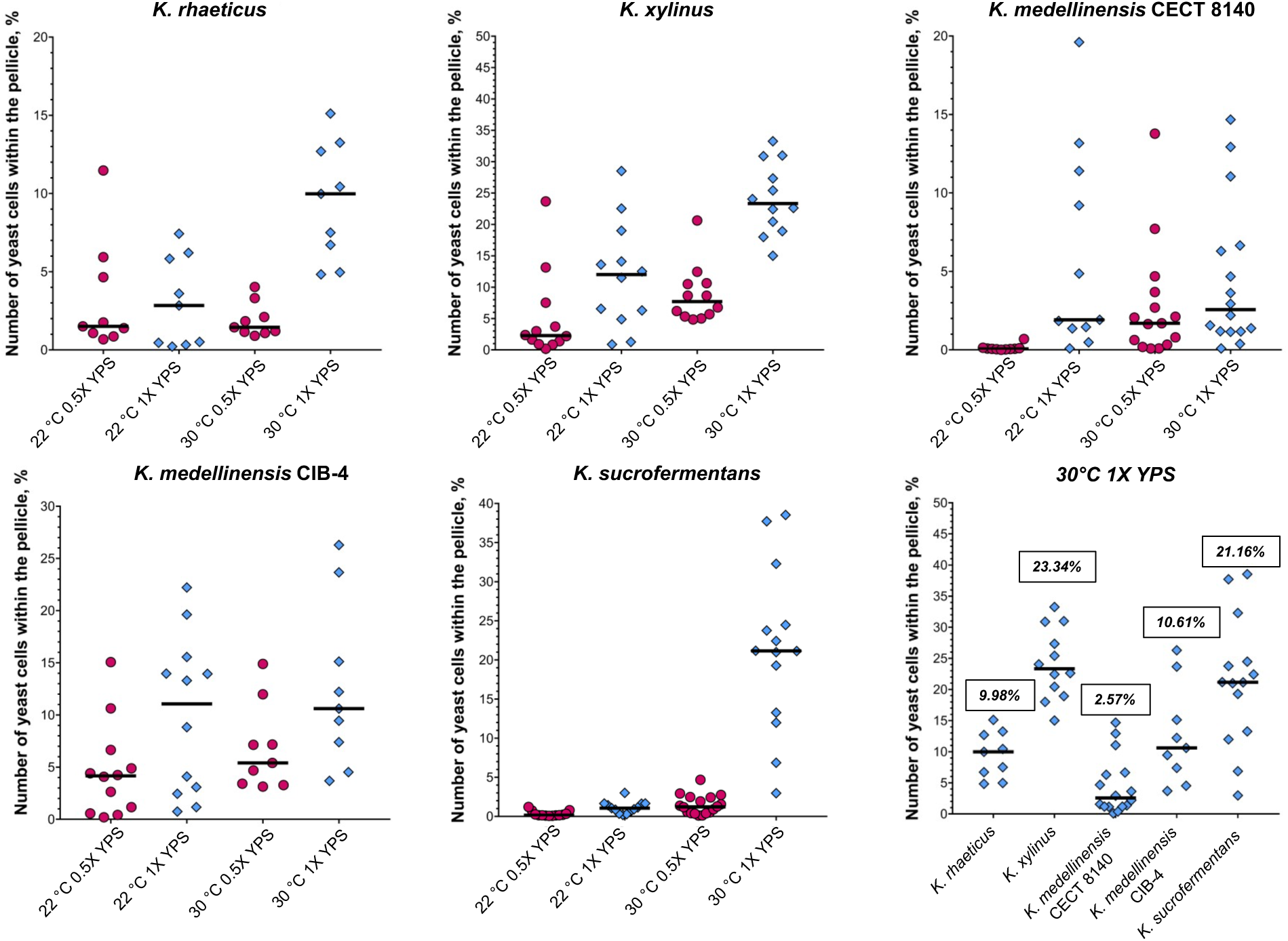
Analysis of yeast incorporation into pellicles. SynSCOBY pellicles were grown at room temperature (22 °C) and 30 °C for 5 days in 0.5X and 1X final concentrations of YPS, digested in PBS+2% cellulase and 2 ml samples were analysed by flow cytometry. Based on FSC-H and SSC-H cells were divided into bacteria and yeast subpopulations. Number of yeast cells was calculated as % from both bacteria and yeast events within the analysed sample. Data was processed with GraphPad Prism (significant outliers removed). Median values for pellicles grown at 30 °C in 1X YPS are displayed on the graph.

Flow cytometry revealed strain-specific differences in yeast incorporation across the four tested conditions. Higher yeast populations were consistently observed in 1X YPS compared to 0.5X YPS at both temperatures. Given that 1X YPS at 30 °C maximised yeast incorporation, these conditions are optimal for SynSCOBY applications. Under these conditions, *K. xylinus* and *K. sucrofermentans* pellicles exhibited median yeast incorporation of 28.76% and 21.16%, respectively. *K. rhaeticus* and *K. medellinensis* CIB-4 showed 9.98% and 10.61%, whereas *K. medellinensis* CECT 8140 demonstrated substantially lower incorporation at 2.57%. Overall, all tested *Komagataeibacter* strains successfully formed functional SynSCOBYs and produced BC-yeast pellicles.

Engineering yeast to produce functional proteins within SynSCOBYs represents an additional BC functionalization strategy. In co-cultures developed with *K. rhaeticus* iGEM, yeast secreted the enzyme beta-lactamase (BLA) ^37^. Beta-lactamase hydrolyses the chromogenic substrate nitrocefin, producing a colour transition from yellow to red that enables visual detection of enzyme activity. We employed an optimised BLA-expressing yeast strain exhibiting high secretion levels of the enzyme^53^. Beta-lactamase was genetically fused to a CBMcex to facilitate enzyme immobilisation within the cellulose matrix. After 5 days of cultivation of bacteria and the engineered yeast, pellicles were washed and nitrocefin was applied topically and colour images were taken. Based on flow cytometry data demonstrating yeast incorporation variability, we anticipated heterogeneity among pellicles produced by individual strains.

Visual inspection of yeast-BLA-CBM pellicles confirmed the expected variation within replicate sets (Fig. 7, left column). Quantitative colorimetric analysis was not performed due to non-uniform spatial coloration. However, inter-strain differences were readily apparent by visual assessment. *K. rhaeticus* (triplicate set) and *K. medellinensis* CIB-4 (one replicate) exhibited deep red coloration, while *K. medellinensis* CECT 8140 (triplicate set) displayed an orange-red coloration, confirming less enzyme presence within pellicles. This result aligns with flow cytometry data showing *K. medellinensis* CECT 8140 giving SynSCOBYs with a lower yeast population (Fig. 6). Surprisingly, both *K. xylinus* and *K. sucrofermentans* exhibited only faint orange-red coloration, despite these two strains demonstrating the highest yeast incorporation levels by flow cytometry.

**Figure 7.**
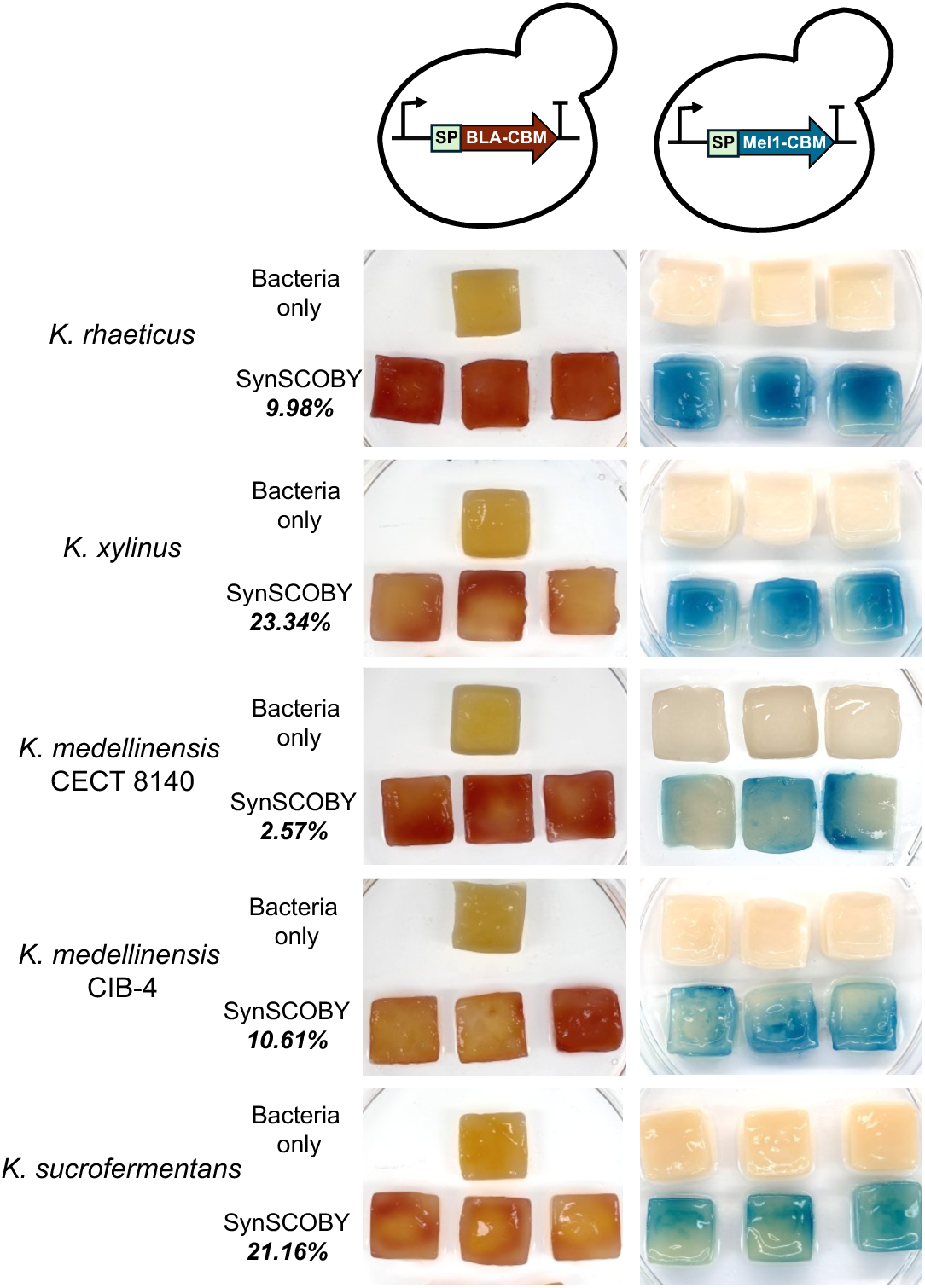
Functionalisation of bacterial cellulose via SynSCOBY. Cellulose-producing bacteria were co-cultured with yeast secreting beta-lactamase (BLA) and alpha-galactosidase (Mel1) fused to cellulose binding motifs. SynSCOBYs were set up in 1X YPS final as described in methods and allowed to form pellicle at 30°C for 5 days. Pellicles were washed in PBS (BLA) and citric buffer (Mel1) prior to reaction. Nitrocefin and X-α-Gal were added topically. Images were taken after 1 hour (BLA) and 16 hours (Mel1) after start of the reaction. Pellicles produced by bacteria monocultures were used as control. Median % of yeast cells incorporated into similar SynSCOBYs established by flow cytometry previously are displayed.

Two hypotheses could explain this discrepancy. First, yeast cells may be metabolically inactive within these SynSCOBYs, preventing beta-lactamase secretion. Second, pellicle microenvironmental conditions (pH, bacterial metabolites) may inhibit the nitrocefin reaction. To address the first hypothesis, we tested SynSCOBYs with yeast secreting an alternative enzyme, alpha-galactosidase (yeast-Mel1-CBM) ^37,53^ (Fig. 7, right column). Mel1 hydrolyses the chromogenic substrate X-α-Gal, producing a colourless-to-blue transition. Mel1 activity was detected in all pellicles. Visual inspection revealed reduced blue coloration in *K. medellinensis* CECT 8140 pellicles, consistent with low yeast cell numbers. Conversely, *K. rhaeticus* pellicles displayed blue intensity comparable to *K. xylinus*, suggesting the reaction in these pellicles may be substrate-limited rather than proportional to number of yeast cells within the pellicle.

Since X-α-Gal conversion was observed in all pellicles, yeast cells must remain metabolically active and capable of enzyme secretion across all SynSCOBYs. We therefore investigated whether *K. xylinus* and *K. sucrofermentans* culture media contain metabolites that inhibit the nitrocefin reaction or whether media pH is restrictive for enzyme activity. We first examined spent media collected from liquid cultures (**Fig. S4A**). Nitrocefin was added to spent media from yeast-BLA-CBM cultures, *K. rhaeticus*, *K. xylinus*, and *K. sucrofermentans*. The latter three samples were supplemented with commercial beta-lactamase. YPD media and YPD supplemented with beta-lactamase served as negative and positive controls, respectively. The most pronounced colour change occurred in yeast spent media. Nevertheless, colour transitions (yellow to orange) were observed in spent media from all BC-producing bacteria with *K. rhaeticus* and *K. xylinus* spent media developing comparable orange colour. While bacterial spent media did not support maximal beta-lactamase activity, enzymatic conversion remained detectable.

We subsequently cultured pellicles in the presence of yeast-BLA-CBM spent media (**Fig. S4B**). Although red coloration was less intense than in SynSCOBY pellicles, pellicles from all three bacteria strains showed comparable beta-lactamase incorporation levels. We also examined whether SynSCOBY-specific conditions for *K. xylinus* and *K. sucrofermentans* inhibit the beta-lactamase reaction, but this was not observed - colour change in spent media was detected across three independent pellicle replicates for both bacterial strains (**Fig. S4C**). Thus, while pH or metabolites in bacterial and SynSCOBY spent media are suboptimal for maximal beta-lactamase activity, media composition alone cannot explain why *K. xylinus* and *K. sucrofermentans* SynSCOBY pellicles reproducibly exhibit less red coloration than anticipated based on flow cytometry results (**Fig. S4D**). Further investigation is required to elucidate the mechanistic basis of this phenomenon.

This work demonstrates that all five BC-producing strains can form functional SynSCOBYs, and these pellicles can be functionalized with yeast-produced enzymes to varying degrees. While flow cytometry enabled quantification of relative cellular composition within pellicles, determining the absolute amount of enzyme secreted by yeast cells in SynSCOBYs and the quantity incorporated into the cellulose matrix remains challenging. We also assessed identical pellicles for cell count and beta-lactamase activity in standard and buffered media (**Fig. S5**). Although all pellicles developed bright red coloration under all conditions, we did not observe a straight correlation between yeast cell numbers within analysed pellicles and chromogenic reaction intensity from enzymatic activity and more investigation is required to address this yeast productivity within SynSCOBY.

### Using biogenic welding for producing multicoloured ELMs

Finally, after exploring two key properties that justify the use BC for ELMs (genetic programmability and functionalization via microbial co-cultures) we sought to demonstrate another of its living material properties: *biogenic welding*. The ability of cut pieces of the living BC material to reconnect by producing new material at their break points inspired us to fabricate composite materials from different engineered bacteria producing distinct chromoprotein outputs ^54^. As a proof of concept, we used *K. rhaeticus* as used in the prior study. We grew independent square pellicles from *K. rhaeticus* monocultures with no genetic changes, or with plasmids added for constitutive expression of AmilCP, or PurpleCP (as described in Fig. 1C). Assorted pellicles were then arranged together in a single culture plate, submerged in minimal media volume, and incubated at 30 °C to enable biogenic welding to seal the gaps between the patches (Fig. 8). The resulting material formed a single contiguous piece comprising twelve differentially coloured pellicles that could be dried as an integrated unit. Although chromoprotein colours faded during drying, distinct chromatic gradations remained detectable in the final material. Following, the approach demonstrated here, one could envision that combining BC sections from bacteria engineered with a variety of inducible genetic constructs in a single assembly plate would further expand the functional design space for ELMs.

**Figure 8.**
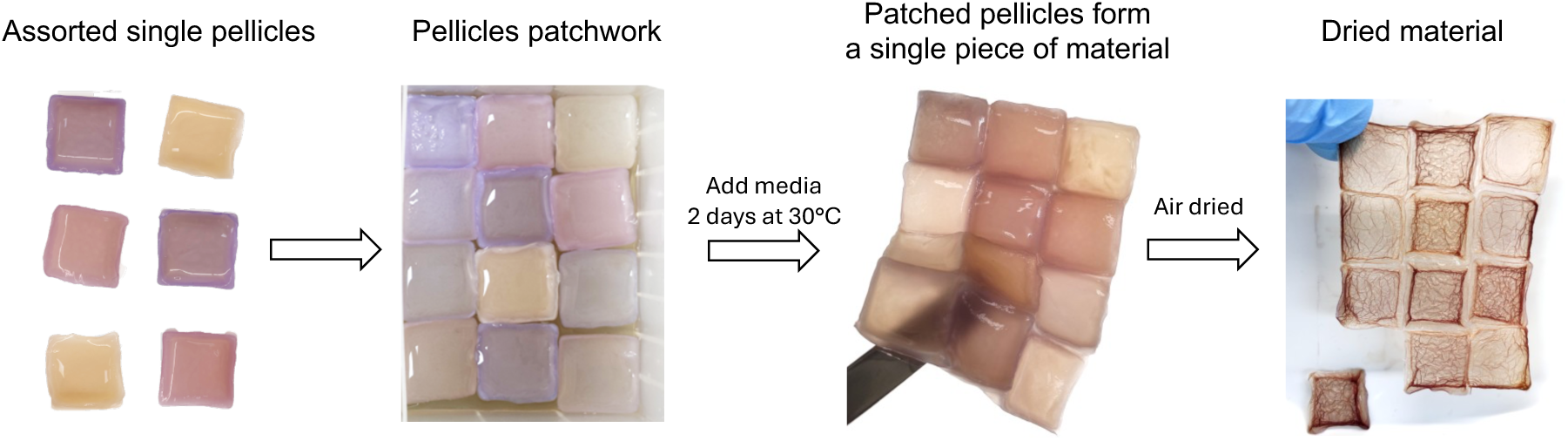
Fabrication of a patchwork material by biogenic welding. Pellicles grown from wildtype *K. rhaeticus* and engineered *K. rhaeticus* strains expressing AmilCP and PurpleCP were grown at 30 °C. Assorted pellicles were then taken and placed next to each other in a single culture plate and covered with YPD. Pellicles were left to weld together for 4 days at 30 °C. The patchwork material was then removed from the container and allowed to air dry for 3 days.

## Discussion

Bacterial cellulose offers potential as a sustainable alternative to some of the mostly widely used materials including plastics and leather ^2–4,6–13^. This study demonstrates the versatility of BC as an ELM extends beyond a single strain, with *Komagataeibacter* species assessed in this work showing distinct advantages for different applications. Our overall findings are summarised in a heat map format (Fig. 9) to guide selection of BC-producing strains when designing for ELM applications.

**Figure 9.**
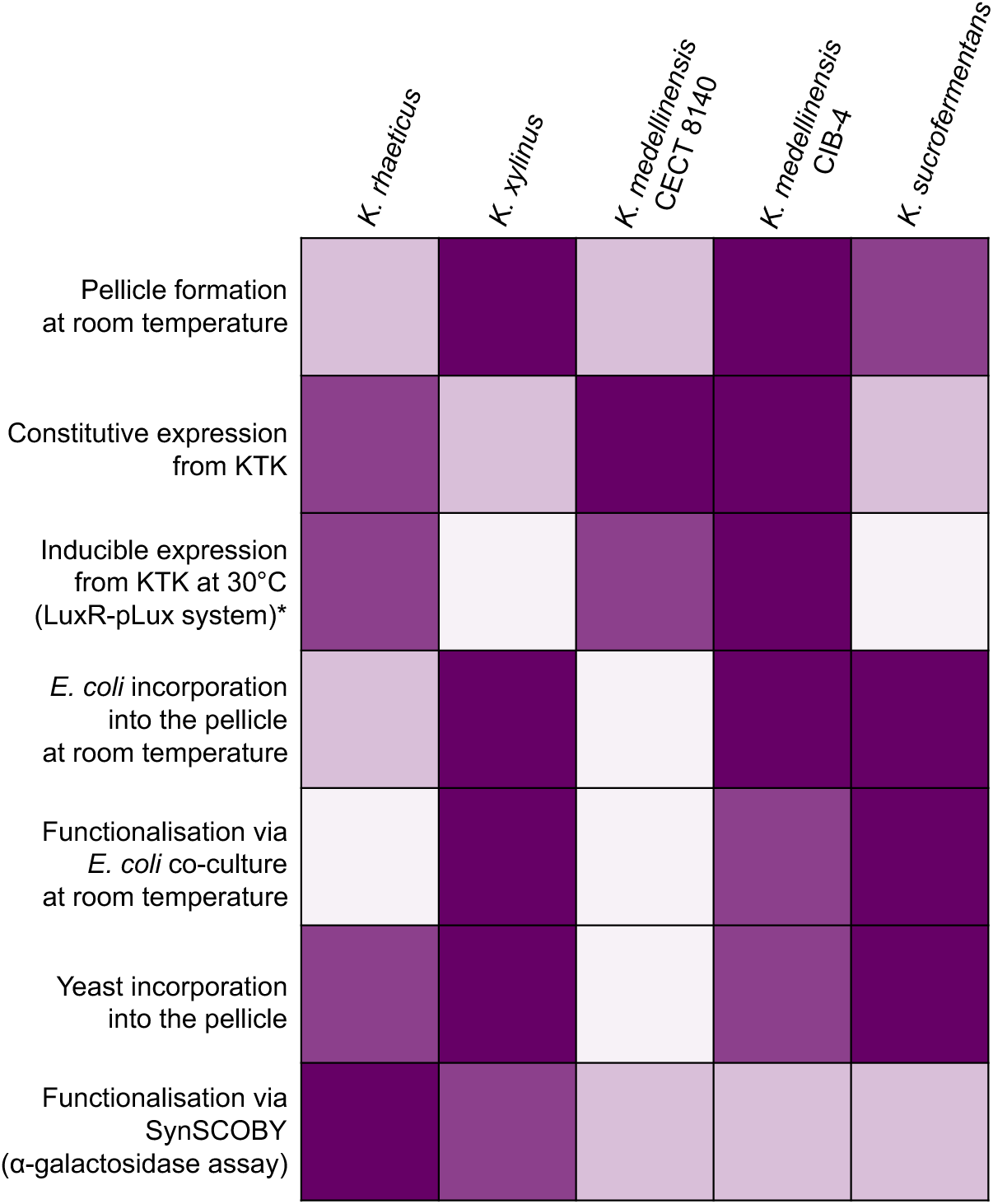
Comparison of performance of different *Komagataeibacter* species. Heatmap based on arbitrary units assigned to each BC-producing bacterium for each specific criterium. Dark shades represent the best performance. (*) Note that with *K. sucrofermentans* performance of pLux-directed gene expression is different at room temperature, and pellicle integrity is affected by AHL addition.

The successful transformation and expression of KTK-based plasmids in all tested strains confirms the broad applicability of this modular cloning system. Our systematic comparison of four widely used species reveals that the KTK toolkit, originally developed for *K. rhaeticus* iGEM ^23^, can be successfully applied across the genus, although with notable variations in performance. *K. medellinensis* strains demonstrated the highest levels of heterologous gene expression, whereas *K. xylinus* and *K. sucrofermentans* showed attenuated mScarlet expression, indicating these strains may require gene expression optimisation, for example by using species-specific promoter and ribosome binding site variants. The KTK system also includes a plasmid for chromosomal integration at the arsenic resistance (ARS) locus. However, the genomic context surrounding the ARS locus varies among tested strains, and the locus is entirely absent in K. *medellinensis*, preventing use of the KTK integration plasmid in these species.

Notably, the failure to see AHL-inducible heterologous expression in *K. xylinus* represents an important limitation for applications using quorum sensing agents. While *K. rhaeticus* and *K. medellinensis* showed clear AmilCP induction with visible colour changes, the absence of response in *K. xylinus* suggests fundamental differences in either LuxR expression, AHL uptake, or downstream transcriptional machinery.

Our co-culture experiments also reveal that the ability to incorporate *E. coli* into growing bacterial cellulose at room temperature is not universal across *Komagataeibacter* species. The striking inconsistency in *E. coli* incorporation by *K. rhaeticus*, ranging from complete absence to 25-30% of the population, stands in sharp contrast to the reproducible co-cultures achieved with *K. xylinus*, *K. medellinensis* CIB-4, and *K. sucrofermentans*. This variability appears linked to pellicle formation efficiency at room temperature, where *K. rhaeticus* and *K. medellinensis* CECT8140 produced noticeably thinner pellicles. The temperature dependency suggests that optimal cellulose production conditions may not align with optimal co-culture conditions, presenting a trade-off that must be considered in experimental design.

All tested strains successfully formed SynSCOBY with *S. cerevisiae*, although yeast incorporation varied substantially. The use of 1x YPS media and cultivation at 30 °C maximised yeast incorporation across most strains, with *K. xylinus* and *K. sucrofermentans* achieving the highest levels. However, high levels of yeast incorporation do not necessarily translate to high levels of enzymatic activity in pellicles where the enzyme is provided by the yeast. While *K. xylinus* may be optimal for applications requiring high yeast incorporation and Mel1 enzyme activity, *K. rhaeticus* appears better suited for producing materials with β-lactamase. This result suggests that substrate- and enzyme-specific compatibility as well as pH compatibility may need to be empirically determined for each desired functionality.

The successful creation of multicoloured composite materials through the biogenic welding properties of BC demonstrates an additional dimension of programmability. By allowing independently grown, functionally distinct pellicles to fuse into a single coherent material, we show that spatial patterning of functionality can be achieved via two-step biosynthesis. While the chromoprotein colours faded upon drying, likely due to protein denaturation or oxidation, this proof-of-concept shows exciting possibilities for creating materials with spatially defined properties. The approach could be extended beyond visual markers to create materials with regions of different enzymatic activities, mechanical properties, or responsiveness to environmental signals. Future work incorporating inducible genetic circuits in different patches could enable materials that respond differentially to inputs depending on spatial location.

To summarise, based on our comprehensive analysis, we propose the following guidelines for strain selection.

- *K. medellinensis* and *K. rhaeticus* are the best species for heterologous gene expression, both for constitutive expression and AHL-induced expression. These strains are optimal for materials requiring dynamic programmability.
- *K. xylinus*, *K. medellinensis* CIB-4, and *K. sucrofermentans* all enable reproducible co-cultures with *E. coli*, with the latter showing the most even incorporation. *K. rhaeticus* and *K. medellinensis* CECT 8140 do not support *E. coli* co-cultures at room temperature.
- *K. xylinus* and *K. sucrofermentans* incorporate the most yeast cells in SynSCOBY pellicles. However, robust functionality of SynSCOBY may need to be established independently for each engineered yeast.
- *K. xylinus* and *K. medellinensis* CIB-4 produce robust pellicles at 22 °C, making them suitable for applications incompatible with 30 °C cultivation, such as temperature-sensitive co-culture or bioprocessing equipment.

While this study provides initial guidance for strain selection, future work should focus on understanding the mechanistic basis for strain differences, particularly the factors controlling co-culture compatibility, inducible system responsiveness, and enzyme activity in pellicles. The substantial batch-to-batch variation observed in both *E. coli* co-culture work and SynSCOBY formation highlights the need for better standardisation of culture conditions, media composition, nutrient availability, cells metabolism and quality control measures. Understanding the sources of this variation, whether from inoculation ratios, minor environmental fluctuations, or stochastic population dynamics, represents an important area for future investigation.

## Materials and Methods

### Plasmid construction and transformation

Strains and plasmids used on the study are listed in Supplementary Tables 1 and 2. Plasmids for expression in *E. coli* were constructed using standard digestion-ligation method. The DNA encoding AmilCP, RedCP-CBM, sfGFP and sfGFP-CBM was synthesized and cloned into pET28a or pET29a via use of the plasmid MCS. Plasmids were transformed into the *E. coli* BL21 (DE3) strain and transformants were selected on LB agar plates supplemented with ampicillin. Plasmids for expression in *Komagataeibacter* (AmilCP-constitutive expression KTK_448 and AmilCP AHL induced expression KTK_449) were constructed using standard Golden Gate cloning methods with KTK parts developed and described elsewhere ^23^. AmilCP DNA was amplified from pET28a_AmilCP-CBM. The plasmids were first transformed into *E. coli* competent cells. Resulting plasmids were then transformed into electrocompetent *Komagataeibacter* cells via electroporation ^20^. Transformants were selected on YPD agar plates (1% Yeast extract, 2% Soybean peptone, 2% Glucose, 2% agar) supplemented with 340 µg/ml chloramphenicol or 100 µg/mL spectinomycin.

### Growing monocultures and pellicles from monocultures

*Komagataeibacter* strains were grown in liquid YPD and 2% cellulase (Sigma, C2730) 3-4 days to high density at 30 °C shaking. Media was supplemented with 340 µg/mL Chloramphenicol or 100 µg/ml Spectinomycin when needed. Cells were washed with fresh YPD, prepared to OD_600_ 2.5 and 50 µl were inoculated into 5 ml of YPD + antibiotics. The cultures were incubated at 30°C static for 5 days to allow pellicle formation. Resulting pellicles were washed in PBS for 1 hour and then imaged.

To assess induction of AmilCP in cellulose producing bacteria cells wt and pLux-AmilCP were grown at 30 °C shaking in 3 ml of YPD supplemented with 2% cellulase and 100 µg/ml Spectinomycin. After reaching turbidity (3 days growth) cultures were divided into two tubes and topped up with fresh YPD/Cellulase and Spectinomycin for pLux-AmilCP cells. AHL (*N*-Acyl homoserine lactone, Sigma, K3255) at concentration of 1 µM was added to one set of tubes. Cultures were allowed to grow for 3 more days at 30 °C shaking. Pellicles were set up as described above. On day 3 of pellicle growth AHL was added at final concentration 1 µM. The pellicle was left to grow for 3 more days, washed in PBS and imaged.

### Co-culturing with *Escherichia coli*

To analyse presence of *E. coli* within pellicles, co-cultures with wildtype *Komagataeibacter* and *E. coli* transformed with pET28-GFP were established. *Komagataeibacter* strains were grown in liquid YPD and 2% cellulase (Sigma, C2730) 3-4 days to high density at 30 °C with shaking. *E. coli* was grown overnight in the presence of ampicillin at 37 °C with shaking. 5 mL of YPD media was aliquoted into 12-well plates and the starter cultures of cellulose producing bacteria and *E. coli* were added in equal volume to give a final ratio of 1:1000 cells (Komagataeibacter: *E.coli*). Cell 10^x^ of both bacteria were calculated using the following formula. The formula was derived from CFU.

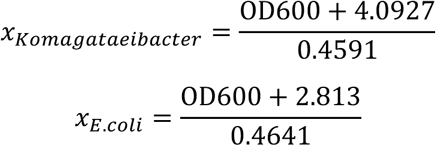

Pellicles were grown statically at room temperature (RT, 22 °C ± 2) for two weeks. To produce coloured BC pellicles, *E. coli* strains carrying pET28_AmilCP and pET28_RedCP-CBM were co-cultures with *Komagataeibacter*. Expression of recombinant proteins was induced with 1 mM IPTG after 24 hrs of co-culturing. Control conditions included YPD without *E. coli* or without cellulose producing strains. Once grown, pellicles were harvested and washed in 10 ml PBS with agitation for 1 hr at RT. Pellicles were then collected at stored at 4 °C for 4 days to allow protein maturation and colour development to occur before imaging.

A sender *E. coli* strain which recombinantly produces AHL was employed to produce another set of coloured pellicles. This *E. coli* was co-cultured with *Komagataeibacter* as described above. Controls for this experiment included YPD without *E. coli* or without cellulose producers, and 1 µM AHL (Sigma, K3255), in place of *E. coli*.

### SynSCOBY and enzymatic reactions with pellicles

For co-cultures, *Komagataeibacter* were grown in YPD with 2% cellulase (Sigma, C2730) 3-4 days to high density at 30 °C with shaking. Yeast strains were grown in YPD overnight at 30 °C with shaking. *Komagataeibacter* cells were washed in fresh YP media and resuspended in YP to OD_600_ = 2.5. Yeast cells were serially diluted to OD_600_ = 0.01 in fresh YP. Co-cultures were set up in 24 deep-well plates with in 6 ml of final volume (45% Optiprep (Sigma, D1556), 1% Ethanol, 1% yeast culture, 2% cellulose producing bacteria and YP-Sucrose to make up 6 ml). The pellicles were grown at static conditions at 30 °C or at room temperature (20 °C ± 2) for 5 days. For pellicle analysis by flow cytometry two different concentrations of YPS were used: standard (1% yeast extract, 2% soybean peptone, 2% Sucrose) and double (2% yeast extract, 4% soybean peptone, 4% Sucrose). Since co-cultures contain 45% of Optiprep, the final concentration of YPS is half of the standard required for optimal microbial growth (0.5X). Therefore, using double of standard concentration leaves us with optimal YPS concentration in the final co-culture (1X). For further cellulose functionalisation via co-cultures with yeast final 1X YPS was used.

*Saccharomyces cerevisiae* strains secreting beta-lactamase-CBM and alpha-galactosidase Mel1-CBM at high levels were engineered in previous work ^37,53^ and SynSCOBY pellicles with these were grown as described above. For the beta-lactamase activity assay, pellicles were washed in PBS for 1 hour and 50 µL of 2 mg/mL nitrocefin in 5% DMSO/95% PBS was then added into the centre of a small pellicle. The pellicles were imaged after 1 hr of reaction. For alpha-galactosidase activity assay pellicles were washed in 100 mM citrate buffer pH 4.5 (45.95 mM sodium citrate dihydrate, 54.05 mM citric acid). 10 µL of 40 mg/mL of X-α-Gal solution in DMSO (Sigma-Aldrich, 16555) were applied into the middle of the pellicle. The reaction was left overnight to develop before imaging.

### Flow cytometry analysis of co-cultures

To analyse cell content in the BC materials, pellicles were grown from co-cultures as described above. For co-cultures with *E. coli*-GFP the pellicles were grown for 14 days at room temperature. Small equal pieces were cut and incubated in PBS with 2% cellulose at 4°C degrees rotating for 48 hours to digest cellulose. The mix was sonicated at 30% working intensity for 20 sec (Vibracell, Sonics&Materials, INC.). Sonication was optimised to separate the cells but to keep them intact (**Fig. S6**). The cell mix was analysed by Attune NxT Flow Cytometer^TM^ (Thermofisher) using Blue Laser 1. The data were analysed using Attune software and GraphPad PRISM. As both bacteria show same forward and side scatters percentage of GFP positive cells was counted as *E. coli* cells. At least 50,000 cells within a “bacteria” gate were analysed for each sample (**Fig. S1**).

For co-cultures with yeast pellicles were grown for 5 days at 30 °C or room temperature. Small equal pieces were cut from the pellicles and incubated in PBS-2% cellulose at room temperature for 48 hours to digest cellulose. The mix was sonicated at 40% intensity for 20 sec (Vibracell, Sonics&Materials, INC). Sonication was optimised to separate the cells but to keep them intact (**Fig. S6**, cells passaged after sonication). The cell mix was analysed by Attune NxT Flow Cytometer^TM^ (Thermofisher). The data were processed with FlowJo (FlowJo, LLC) with appropriate gates applied. Bacteria and yeast shown distinct forward and side scatter characteristics and the number of cells falling into “yeast” gate were calculated as percentage of all events in the “co-culture” gate. At least 20,000 cells within co-culture gate were analysed for each sample.

### Microbial growth curves

*Komagataeibacter* strains were grown in liquid YPD with 2% cellulase (Sigma, C2730) to high density at 30 °C with shaking. Cells were washed in fresh YP and set to OD_600_ = 1.0. 100 µL of cells were put into 1 mL of YPD with 2% cellulase in a 24 well plate. The change in OD_600_ over 100 hours at 22 °C or 30 °C with orbital shaking was measured by a Tecan Spark plate reader. Data were analysed using GraphPad prism and Excel Microsoft. Blank values from media-only wells were subtracted from the main data. Mean values for at least three repeats were used for plots and for generating the standard deviation values.

### Beta-lactamase assay with spent media and pellicles produced by monocultures

For beta-lactamase reaction in the spent media *K. rhaeticus*, *K. xylinus* and *K. sucrofermentans* were grown for 3 days at 30 °C with shaking in YPD with 2% cellulase. Yeast was grown overnight at 30°C with shaking in YPD media. Cultures were centrifuged and supernatant was collected as spent media. 100 μl of supernatant was mixed with 20 μL of 10 μg/mL commercial beta-lactamase (ProspecBio, ENZ-351) and 5 μL of 2 mg/mL nitrocefin. YPD and YPD with added beta-lactamase were used as controls. Reactions were developed for 1 hr prior to imaging the results.

*K. rhaeticus*, *K. xylinus* and *K. sucrofermentans* were grown in YPD with 2% cellulase until dense. Cells were washed in fresh YP and diluted to OD_600_ = 2.5. Pellicles were set up with 50 µL of cells in YPD supplemented with 100 μL of yeast-BLA spent media to assess incorporation of BLA-CBM into the pellicle and were grown for 5 days at 30 °C. Pellicles were washed in PBS for 1 hr and 50 µL of 2 mg/mL nitrocefin in 5% DMSO/95% PBS was added into the centre of each pellicle. The pellicles were then imaged after 1 hr of reaction.

### Patchwork biogenic welding

*K. rhaeticus* wildtype strain and the strains expressing AmilCP and PurpleCP were grown in YPD with 2% cellulase until dense at 30 °C. Cells were washed in fresh YP and diluted to OD_600_ = 2.5. Pellicles were set up with 50 µL of cells in YPD supplemented with 340 µg/mL chloramphenicol for AmilCP and PurpleCP plasmid selection and 100 µL of 100% ethanol for wildtype cells (since chloramphenicol is dissolved in 100% ethanol). Pellicles were grown for 5 days at 30 °C. Formed pellicles were placed into a single plate. Fresh YPD with chloramphenicol was then added into the plate until it just covered the pellicles (5 mL). Patchwork material was harvested after 4 days of incubation at 30 °C and allowed to air dry for 3 days.

## SUPPORTING INFORMATION

A **Supporting Information** file is provided containing supplementary figures, representative raw flow cytometry graphs, microbial growth curves, images of additional BC pellicles, additional data for the beta-lactamase assays, and full descriptions of the strains and plasmids used in the study.

## AUTHOR CONTRIBUTIONS

TE, MDR and MZ developed the concept, AK and KG designed the research, AK, KG and MM performed the experiments, AK, KG and MM analysed the data, AK, KG, MZ and TE wrote the manuscript, AP developed the strain CIB-4 and AP, MZ and MDR all reviewed the manuscript.

## ACKNOWLEDGEMENTS

This work was funded by UKRI Engineering and Physical Sciences Research Council (EPSRC) award EP/Y008243/1 to KG, MDR, MZ and TE. MM is a recipient of an EMBO Scientific Exchange Grant 9225. The strain *K. medellinensis* CIB-4 was generated under the research grant SMARTERIAL PID2023-146557OB-C21 funded by MCIU/AEI/10.13039/501100011033. We would also like to thank everyone who was worked on bacterial cellulose in both Tom Ellis and Meng Zhang’s groups between 2023 and 2026 for their ongoing feedback on the work described here.

## SUPPORTING INFORMATION FILE

**Figure S1.**
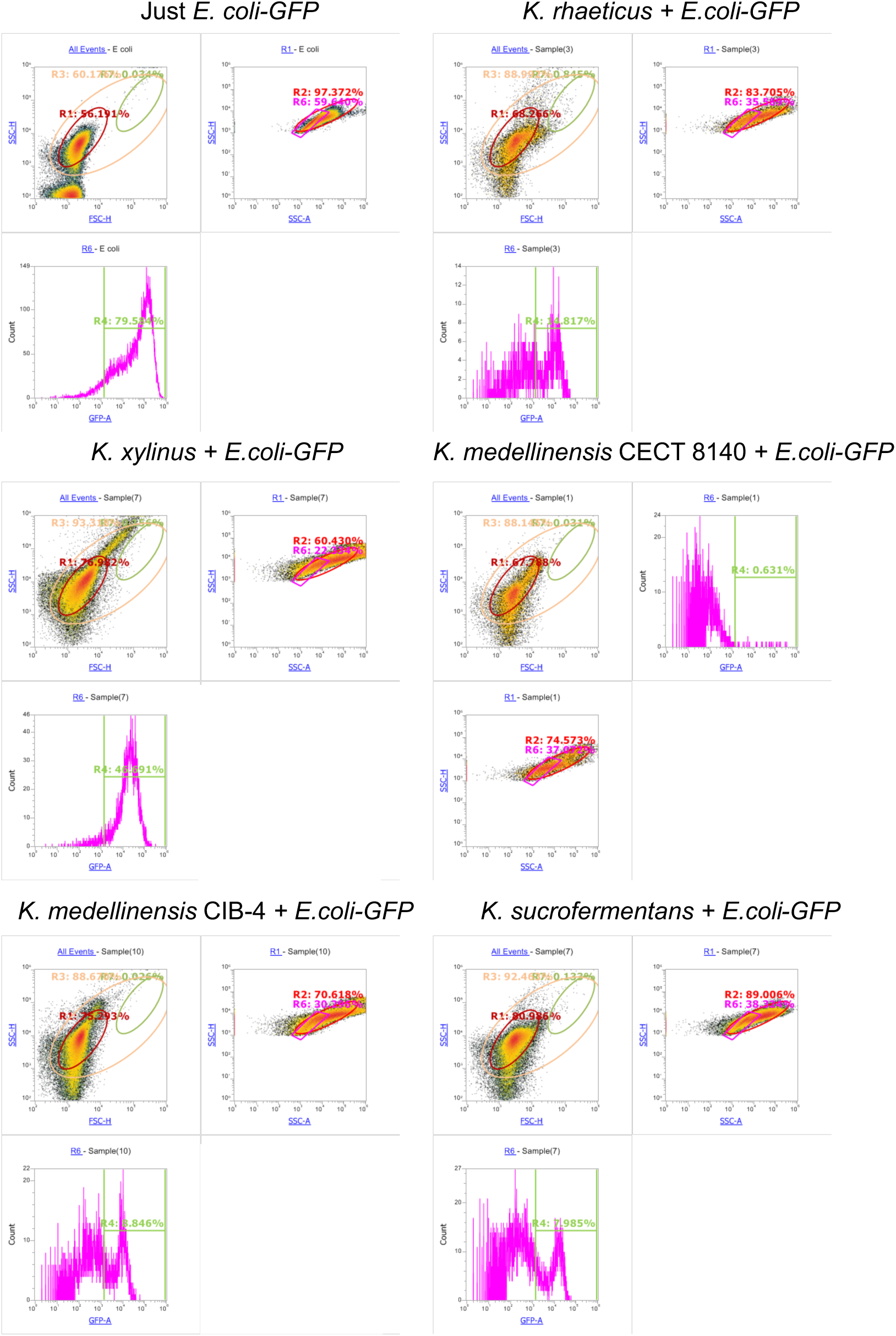
Representative images from flow cytometry analysis of pellicles produced by *E. coli* and cellulose-producing bacteria. Pellicles were grown at room temperature for 14 days, cut and a piece was digested in PBS with 2% cellulase. R1 gate encompasses bacteria cells, R6 – single bacteria cells, R4 - GFP positive cells from R6 gate.

**Figure S2.**
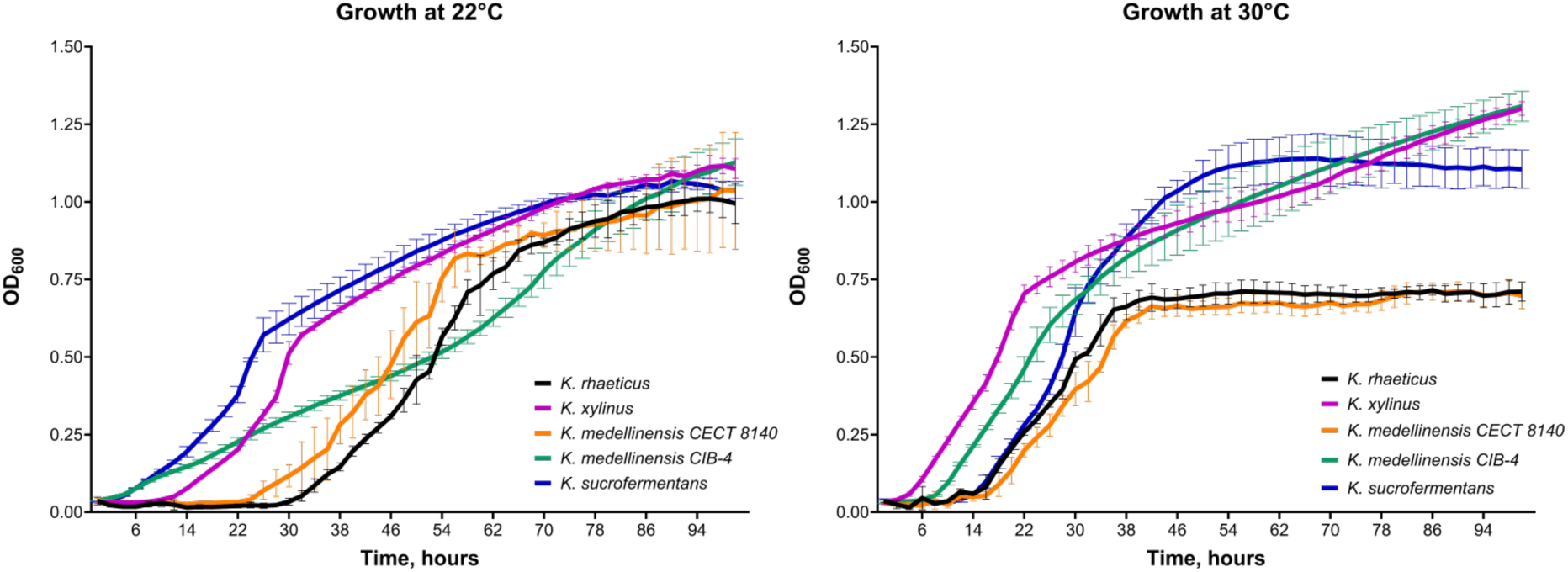
Growth curves of *Komagataeibacter* cells at 22 °C and 30 °C in YPD. Pre-cultures were grown until reaching turbidity (3 days growth) at 30 °C shaking, washed in fresh YPD with 2% cellulase and diluted to OD600 = 0.03 in 1 ml of fresh YPD with 2% cellulase in 24 well plate. Plates were incubated for 100 hours with orbital shaking at required temperature. OD600 was measured every 2 hours. Data was analysed using GraphPad Prism, n = 3 for *K. rhaeticus* and *K. medellinensis* CIB-4 22°C, n = 4 for the rest of the strains, mean value is plotted, error bars – standard deviation.

**Figure S3.**
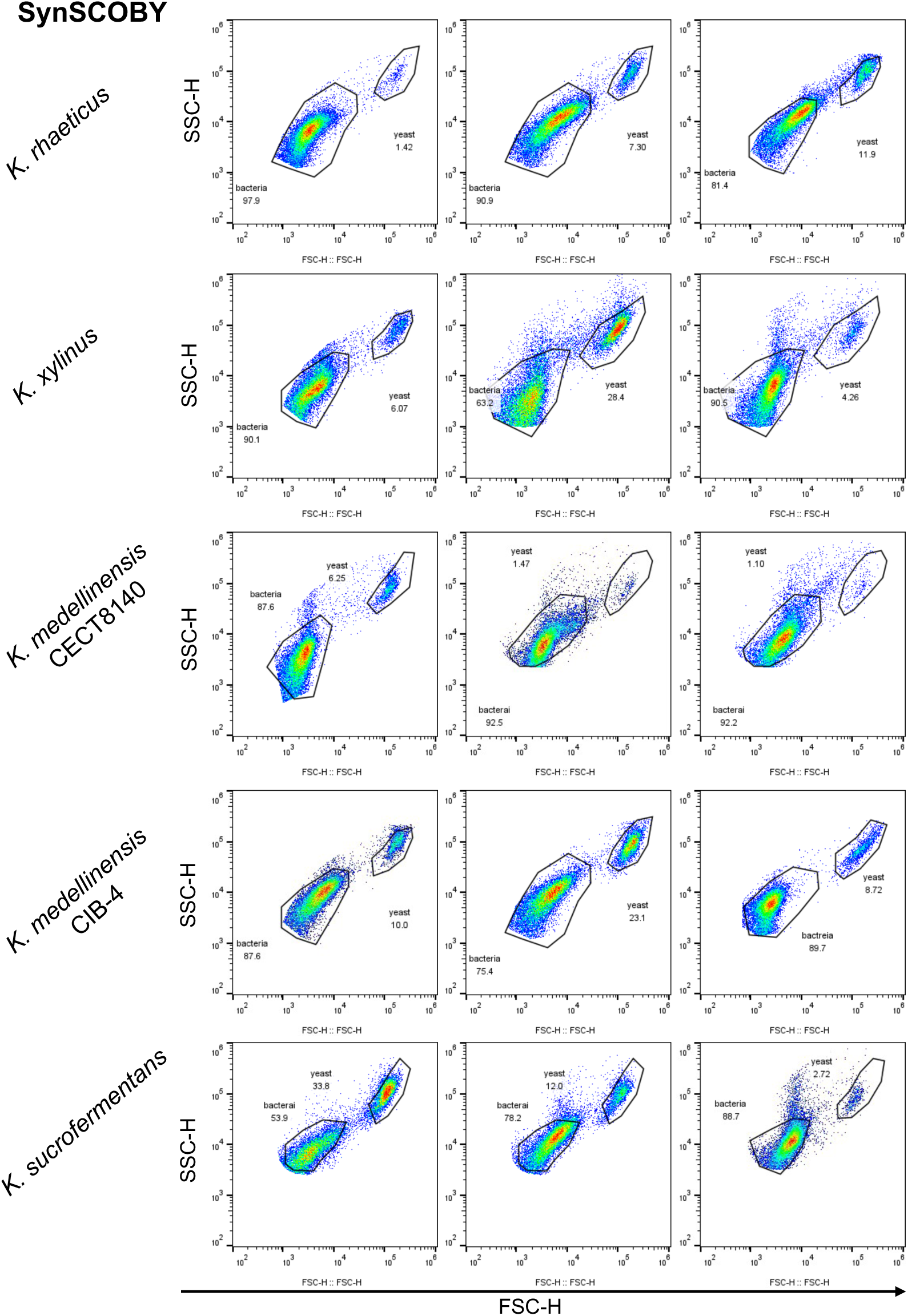
Representative images from flow cytometry analysis of SynSCOBY pellicles. Pellicles grown at 30 °C for 5 days in final concentration of 1X YPS. Pellicles are cut in 4 and 1 piece is digested in PBS with 2% cellulase, sonicated, and 2 mL sample is analysed by flow cytometry.

**Figure S4.**
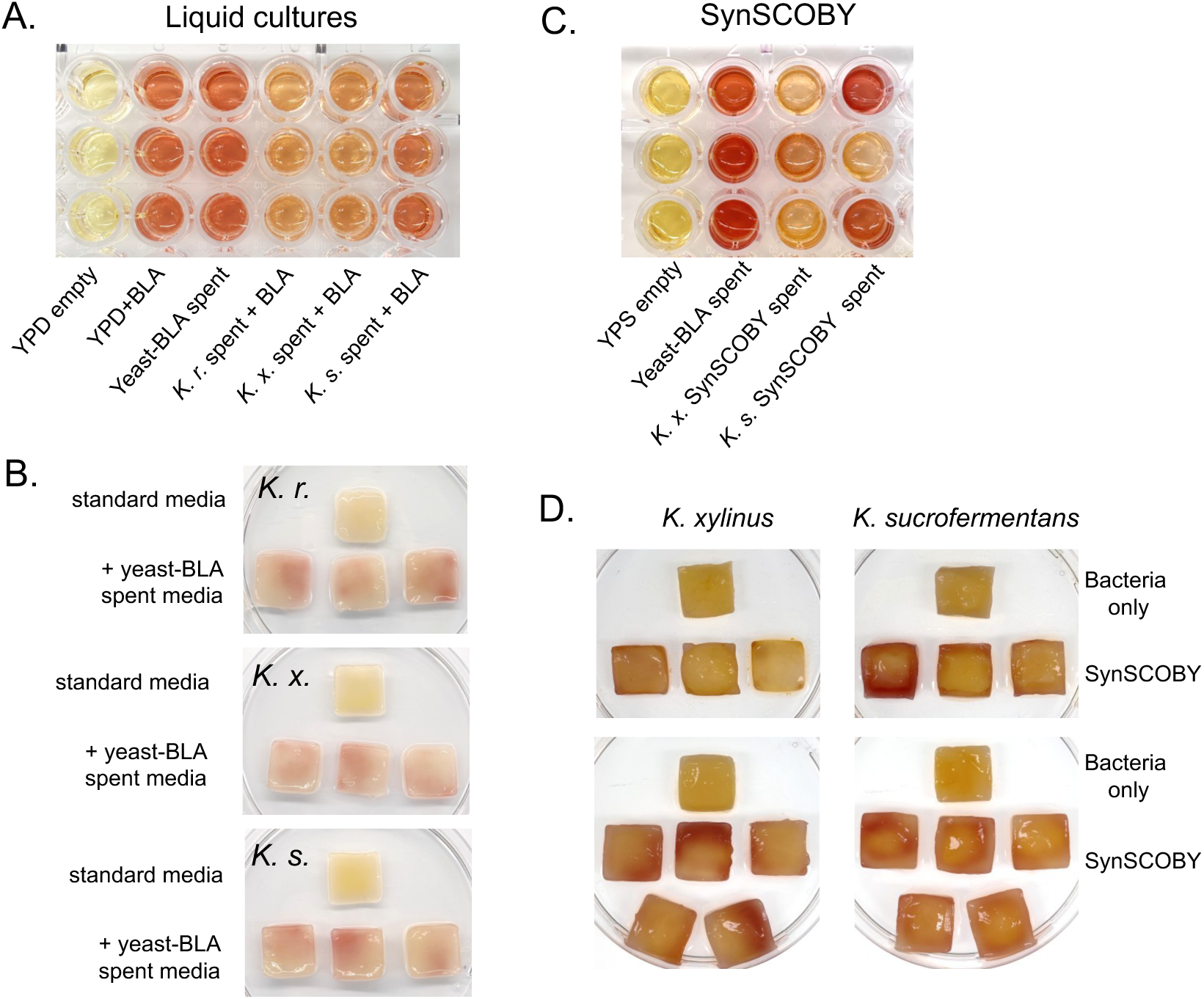
Beta-lactamase activity reaction in spent media and BC pellicles. A) Spent YPD media was collected from yeast secreting beta-lactamase (Yeast-BLA), wt *K. rhaeticus (K.r)*, *K. xylinus (K.x.)* and *K. sucrofermentans (K.s.)*. Media from bacteria was supplemented with commercial beta-lactamase. YPD empty and YPD supplemented with beta-lactamase were used as controls. 5 µl of nitrocefin were added into each well. Technical triplicates were tested. The reaction was run for 1 hour. B) Pellicles from monocultures were grown in YPD supplemented with yeast-BLA spent media containing beta-lactamase-CBM. Pellicles were grown for 5 days at 30 °C. 50 µl of nitrocefin were added to each pellicle. The reaction was run for 1 hour. YPD does not inhibit nitrocefin reaction, and there is no colour transition in empty YPD. Meid was collected from three independent pellicles. The reaction demonstrated variations observed in pellicles. C) Spent YPS media from *K. xylinus* and *K. sucrofermentans* was collected. There is no colour transition in empty YPS (negative control). Yeast-BLA YPD spent media used as a reference for the activity of beta-lactamase secreted by yeast. 5 µL of freshly made nitrocefin were added into each well. Fresh stock of nitrocefin was used what explains more intense colours. The reaction was run for 1 hour. D) SynSCOBYs were grown in 1X YPS final for 5 days at 30 °C. Pellicles were washed in PBS. 50 µL of nitrocefin were added to each pellicle. The reaction was run for 1 hour. Pellicles on the top and bottom panels are from two independent experiments with different precultures.

**Figure S5.**
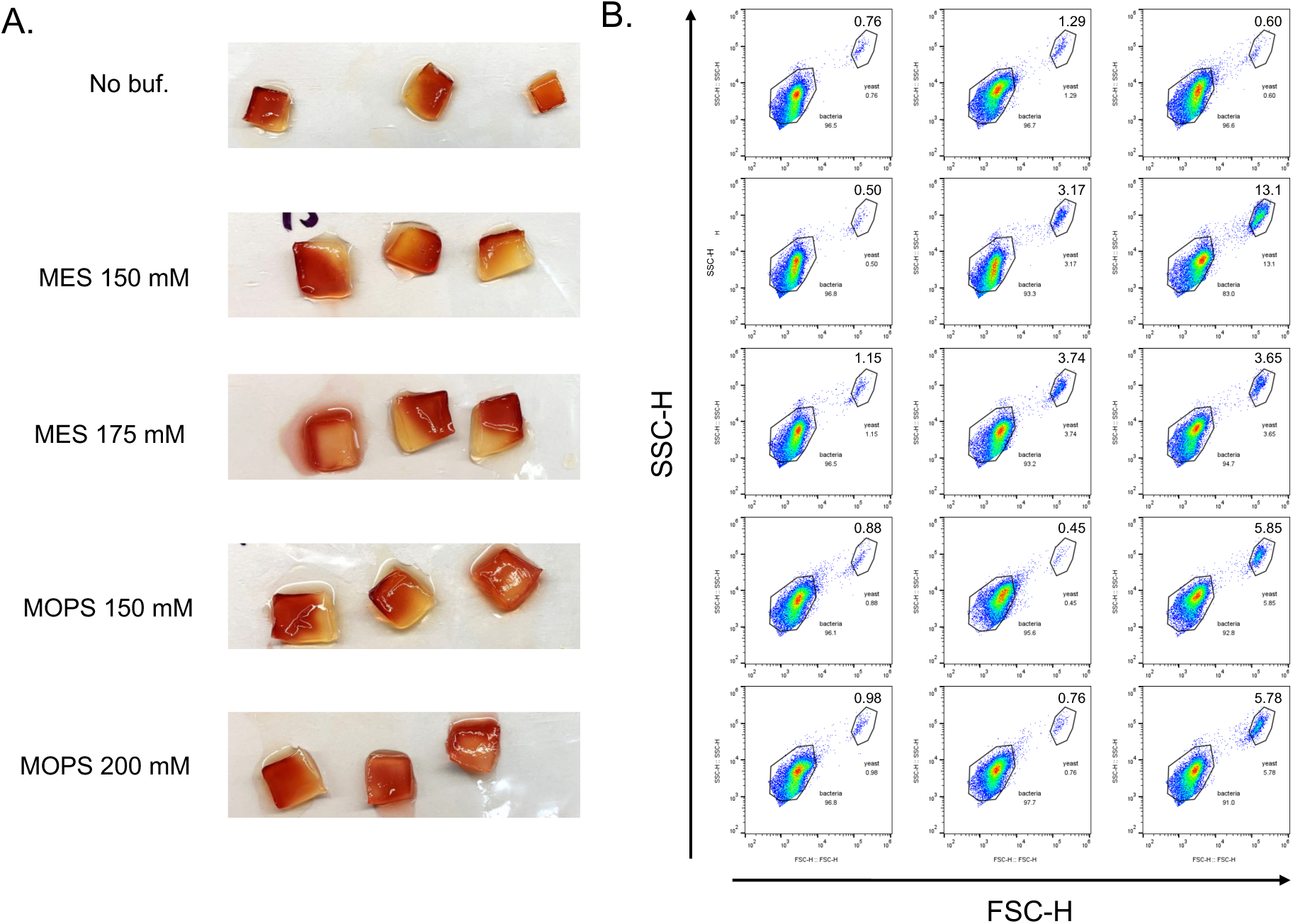
Analysis of *K. rhaeticus* and yeast-BLA SynSCOBY. A) Nitrocefin reaction of SynSCOBY pellicles. Pellicles grown from *K. rhaeticus* and yeast secreting beta-lactamase at 30 °C for 5 days in final concentration of 1x YPS in unbuffered and buffered conditions. After 5 days the pellicle was cut into 4 pieces, and 1 piece was washed in PBS for nitrocefin test. 10 µL of 2.0 mg/mL of nitrocefin in DMSO/PBS was applied onto each and images were taken after 1 hour of reaction. Pellicles were grown from three independent *K. rhaeticus* precultures. B) Flow cytometry analysis of another piece of the same pellicle. Numbers are the percentage of yeast cells out total co-culture gate.

**Figure S6.**
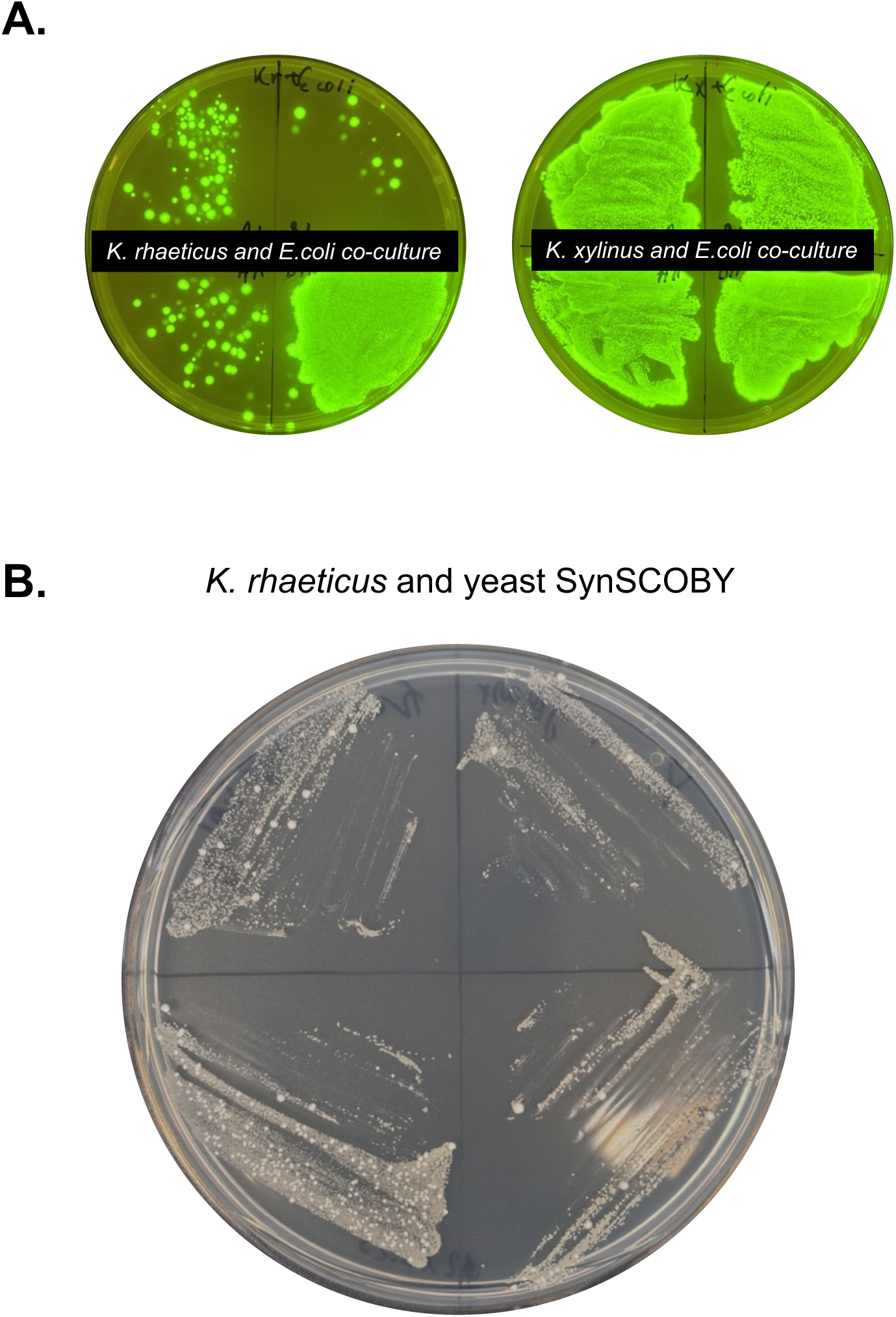
Growth of bacteria and yeast cells after sonication. After sonication some of the sample from digested pellicles were plated onto LB-Amp (A) or synthetic yeast media (B). *E. coli* – GFP are growing and expressing GFP (A), yeast (large white colonies, B) and *K. rhaeticus* (streaks of smaller colonies, B) also growing after sonication.

---

**Supplementary Table 1.**
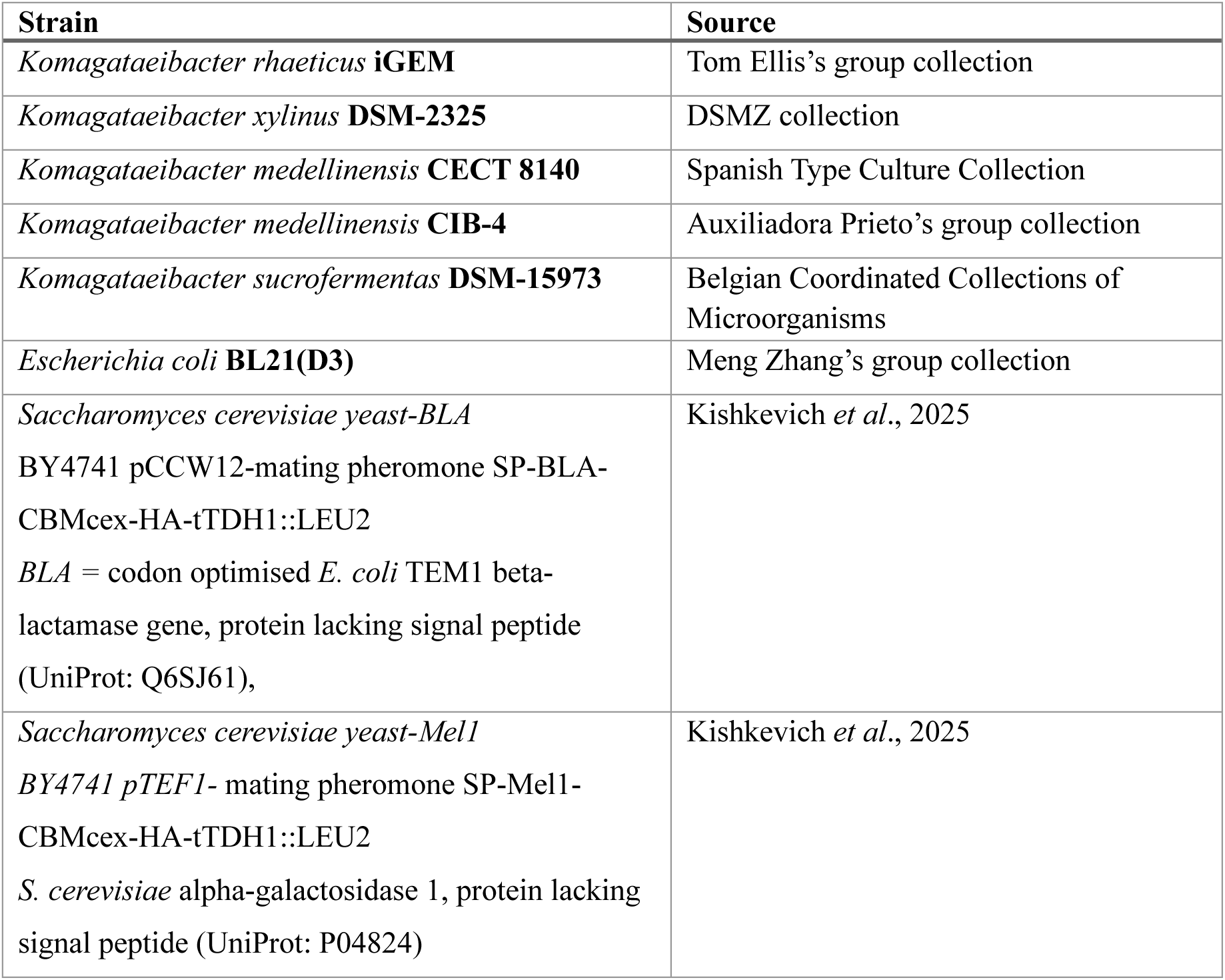
Strains used in this study.

**Supplementary Table 2.** Plasmids used in this study.

| Plasmid name | Construct details | Source |
| --- | --- | --- |
| KTK_349_D1.1_gfasPurple | KTK_136+KTK_002+KTK003+KTK_346+KTK005 | Goosens et al., 2021 |
| KTK_292 | mScarlet plasmid | Goosens et al., 2021 |
| KTK_447_E1.3 AmilCP | KTK_001+AmilCP | This study |
| KTK_448_D1.2_AmilCP constitutive expression | KTK_010+KTK_002+KTK_003+KTK_447+KTK_031 | This study |
| KTK_449_D2.2A_LuxR+pLux AmilCP | KTK_013+KTK_074+KTK_450 | This study |
| KTK_450_D1.2 pLux AmilCP | KTK_010+KTK_034+KTK_003+KTK_447+KTK_031 | This study |
| KTK_074_D1.1 LuxR | KTK_009+KTK_002+KTK_003+KTK_033+KTK_005 | Tom Ellis's group collection |
| pSenderLux01 | CmR, pSEVA331Bb derivative carrying LuxI gene under control of *pJ23104* | Walker et al., 2018 |
| pET29_sfGFP | Constitutive GFP expression | Meng Zhang's group collection |
| pET28a_RedChrome_CEX (RedCP-CBM) | IPTG inducible Red chromoprotein with cellulose binding module | This study |
| pET28a_AmilCP | IPTG inducible Blue chromoprotein | Meng Zhang's group collection |
| pET28a_sfGFP_dCBM (GFP-CBM) | IPTG inducible green fluorescent protein with cellulose binding module | Gilmour et al., 2023 |

## Protein sequences used in this study

### sfGFP

MAHIVMVDAYKPTKGSENLYFQGGGSGRKGEELFTGVVPILVELDGDVNGHKFSVRGEGEG DATNGKLTLKFICTTGKLPVPWPTLVTTLTYGVQCFARYPDHMKQHDFFKSAMPEGYVQERT ISFKDDGTYKTRAEVKFEGDTLVNRIELKGIDFKEDGNILGHKLEYNFNSHNVYITADKQKNG IKANFKIRHNVEDGSVQLADHYQQNTPIGDGPVLLPDNHYLSTQSVLSKDPNEKRDHMVLLE FVTAAGITHGMDELYKGSGSGSHHHHHH*

### Red chromoprotein

MSVIKQVMKTKLHLEGTVNGHDFTIEGKGEGKPYEGLQHMKMTVTKGAPLPFSVHILTPSH MYGSKPFNKYPADIPDYHKQSFPEGMSWERSMIFEDGGVCTASNHSSINLQENCFIYDVKFHG VNLPPDGPVMQKTIAGWEPSVETLYVRDGMLKSDTAMVFKLKGGGHHRVDFKTTYKAKKP VKLPEFHFVEHRLELTKHDKDFTTWDQQEAAEGHFSPLPKALPPTPTPTTPTPTPTTPTPTPTS GPAGCQVLWGVNQWNTGFTANVTVKNTSSAPVDGWTLTFSFPSGQQVTQAWSSTVTQSGSA VTVRNAPWNGSIPAGGTAQFGFNGSHTGTNAAPTAFSLNGTPCTVG*

### Amil chromoprotein

MSVIAKQMTYKVYMSGTVNGHYFEVEGDGKGKPYEGEQTVKLTVTKGGPLPFAWDILSPQC QYGSIPFTKYPEDIPDYVKQSFPEGYTWERIMNFEDGAVCTVSNDSSIQGNCFIYHVKFSGLNF PPNGPVMQKKTQGWEPNTERLFARDGMLLGNNFMALKLEGGGHYLCEFKTTYKAKKPVK MPGYHYVDRKLDVTNHNKDYTSVEQCEISIARKPVVA*

